# Extending the capabilities of deconvolution to provide cell type specific pathway analysis of bulk RNA-seq data for idiopathic pulmonary fibrosis

**DOI:** 10.1101/2024.07.30.605740

**Authors:** Mifflin-Rae Calvero, Sri Harsha Meghadri, Alfonso Carleo, Antje Prasse, David S. DeLuca

## Abstract

**Motivation:** Transcriptome data are confounded by differences in cell type proportions. Differentiating between regulated changes in gene expression and changes due to differing cell type proportions remains a challenge. Therefore, we apply a deconvolution method to correct for changes in cell type proportions and provide a novel cell-type specific pathway analysis.

**Results:** We demonstrate the technique in the context of idiopathic pulmonary fibrosis. Inferred cell type proportions indicated a significant increase in fibroblasts, myofibroblasts, and a decrease in vascular endothelial capillary cells. Pathway analysis after adjustment for proportions indicated IPF-related changes in extracellular matrix organization and TGF-β regulation. Cell-type specific pathway analysis suggested the role of interferon signaling in ATII cells. These results demonstrate that deconvolution is not only useful for assessing cell type proportions, but also can provide cell type-specific pathway analysis, allowing for a much more nuanced interpretation of bulk RNA-seq data.

## 1 Introduction

Prior to the advent and widespread adoption of single cell transcriptomics, bulk RNA sequencing (RNA-seq) was heavily utilized to profile disease conditions. The development of RNA-sequencing has enabled the transcriptomic profiling of biological samples, allowing us to compare between healthy and diseased conditions. Bulk RNA-seq samples contain a mixture of cell types and the expression level of each gene is averaged across these cell types (Kanter and Kalisky 2015; Stegle, Teichmann, and Marioni 2015). Traditional RNA-seq analysis such as differential gene expression analysis does not take into account the cell type composition of the samples, thus confounding the analysis (Cobos et al. 2018). The observed changes in gene expression might be due to the underlying disease status of the samples or due to the differences in the cell type composition of the samples, or a mixture of both (Cobos et al. 2018).

This led to the development of computational approaches, termed deconvolution, to infer the cell type proportions in bulk RNA-seq samples, and it allows for re-analysis of publicly available bulk RNA-seq data sets efficiently (Cobos et al. 2018; Wang et al. 2019). These methods use a range of different statistical models for estimation, from regression-based (Wang et al. 2019; Newman et al. 2019; Jew et al. 2020) to deep learning techniques (Torroja and Sanchez-Cabo 2019; Menden et al. 2020).

In this study, we utilize the disease idiopathic pulmonary fibrosis (IPF) as the research context for applying and developing deconvolution methods. IPF is a chronic and progressive lung disease of unknown cause, which occurs primarily in middle-aged and older persons, where the scarring and stiffening of the lung tissue leads to a decline in lung function. Studies have already used deconvolution in analyzing IPF bulk samples (Ghosh et al. 2022; Huang et al. 2023), however, there is a lack of utilizing the cell type proportion estimates in the downstream analysis. It has been shown that correcting for cell type proportions improved the sensitivity of detecting genes related to a disease (Wang, Master, and Chodosh 2006; Kong et al. 2019; Patrick et al. 2020; Pellegrino-Coppola et al. 2021).

We therefore propose an analysis pipeline, using deconvolution to estimate the cell type proportions in bulk RNA-seq data, and then correcting for cell type variations in the differential expression analysis, and finally performing cell type-specific pathway analysis (Fig. 1). We postulate that this approach would improve the current practice in analyzing bulk RNA-seq data to understand complex biological processes in IPF.

**Figure 1.**
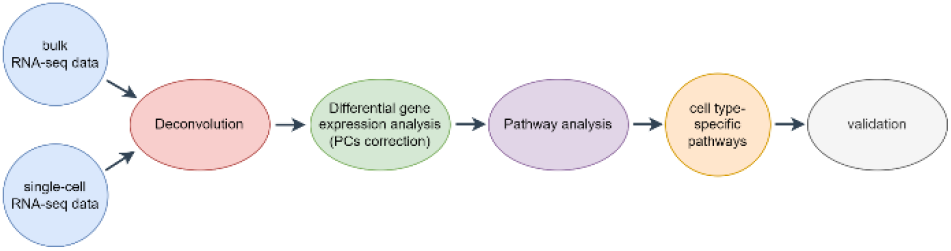
Overview of the analysis procedure.

## 2 Methods

An overview of the analysis procedure is shown in Fig. 1. We analyzed two bulk RNA-seq datasets from GEO as shown in Table 1. All datasets are pre-processed. For bulk RNA-seq data, genes are filtered using the function *FilterByExpr* and then normalized by *calcNormFactors* and *cpm* from the R package *edgeR* (Robinson, McCarthy, and Smyth 2010), and unsupervised analysis of the pre-processed data (Fig. S1-S2).

**Table 1.**
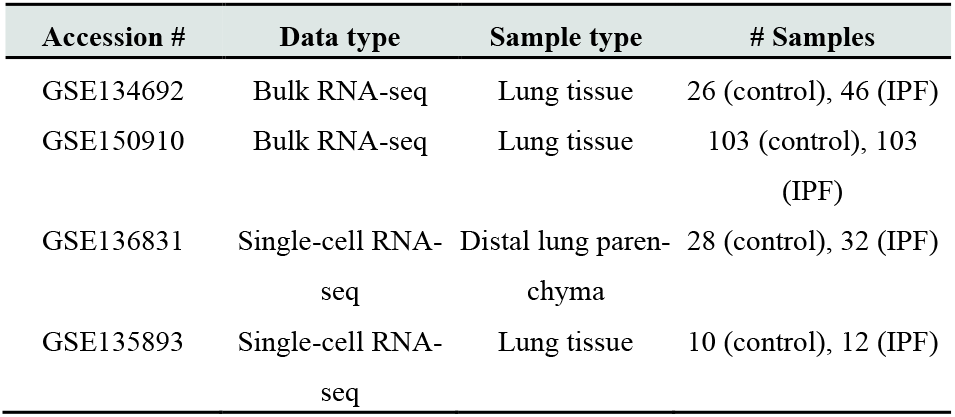
Overview of the datasets.

Differential gene expression analysis of the bulk RNA-seq datasets was performed using the limma-voom approach from the R package *limma* (Ritchie et al. 2015), and a gene was considered to be differentially expressed if the Benjamini-Hochberg adjusted p-value ≤ 0.05. Pathway analysis was then performed using the R package *enrichR* (Chen et al. 2013).

Deconvolution of bulk RNA-seq data was performed using the R package *MuSiC* (Wang et al. 2019). The method takes a bulk RNA-seq data with raw counts and a reference scRNA-seq data as input and utilizes a non-negative weighted least squares (W-NNLS) method for estimation of the cell type proportions. MuSiC’s hierarchical clustering tree-guided procedure to handle closely related cell types to recursively estimate the proportions was utilized. It was run with *music_prop*.*cluster* and the intra-cluster markers as additional input. The differences of estimated cell type proportions between control and IPF were also tested using the Mann-Whitney Test, and adjusting for multiple tests. An adjusted p-value ≤ 0.05 was considered to be significant.

To correct for the cell type proportions in the differential expression analysis, we performed a principal component analysis (PCA) on the estimated cell type proportions to reduce the number of covariates. Cell types with estimated zero proportions for all the samples were not included in the PCA. The top principal components (PC) were included, using the scree plot, as additional covariates to the linear model in the differential expression analysis. An adjusted p-value ≤ 0.05 was considered to be significant.

We also performed cell-type specific pathway analysis using *enrichR*. Cell types with estimated zero proportions for all the samples were not included. First, for each cell type, we calculated the correlation coefficient (*R*) between the cell type proportions and the normalized gene expression (log2CPM). Genes with R ≥ 0.5 were considered cell-type specific and used in the subsequent pathway analysis.

We validated the cell-type specific pathways by doing a per cell type pathway analysis on a different scRNA-seq data set and comparing the resulting pathways with Seurat (Satija et al. 2015; Hao et al. 2021).

## 3 Results

### 3.1 Differential expression analysis without deconvolution

We performed differential expression analysis and pathway analysis of the two bulk RNA-seq datasets (Fig. 2). In the first dataset, we identified 2546 up-regulated and 3166 down-regulated genes in IPF, adjusting for batch, age, gender, and smoking (Fig. 2a). We identified TGF-β regulation of extracellular matrix, cholesterol biosynthesis, signaling by PDGF to be the top three pathways (Fig. 2d). In the second dataset, we identified 4361 up-regulated and 4350 down-regulated genes in IPF, adjusting for sex, age, race, smoking, plate, sample, and institution (Fig. 2a). We identified TGF-β signaling pathway, TGF-β regulation of extracellular matrix and interleukin-2 sign aling pathway to be the top three pathways (Fig. 2e).

**Figure 2.**
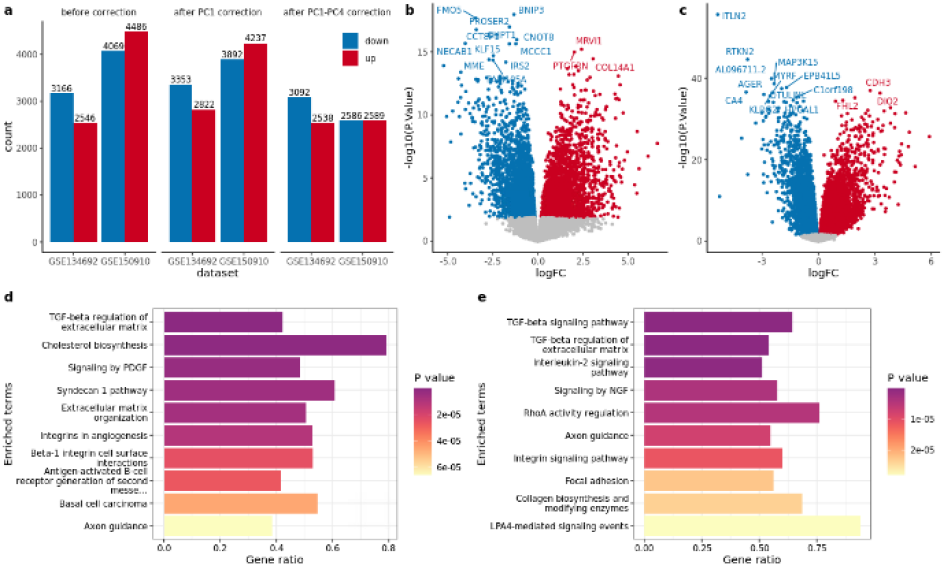
Differential expression and pathway analysis of the two datasets. (a) Number of differentially expressed genes, (b) volcano plot for GSE134692 and (c) GSE150910, (d) top 10 pathways for GSE134692 and (e) GSE150910.

### 3.2 Deconvolution to estimate cell type proportions

Next, we performed deconvolution of the bulk RNA-seq datasets and tested for the difference between the estimated cell type proportions of control and IPF samples. For the first dataset, we identified in IPF a significant decrease in the estimated proportions of alveolar epithelial type 1 (ATI) and type 2 (ATII) cells, vascular endothelial (VE) capillary cells, while a significant increase of smooth muscle cells, pericytes, fibroblasts, myofibroblasts, mesothelial and ciliated cells, and cytotoxic T cell and plasma B cells. Basal, goblet, club cells, aberrant basaloid and T cells were also significantly increased in IPF, although in very small proportions. ILCs, mast cells, B cells, regulatory T cells, cDC1, and mature DCs were estimated to be zero (Fig. 3a).

For the second dataset, we found that the estimated proportions of ATI, VE capillary cells, lymphatic, classical and non-classical monocytes were significantly decreased in IPF, while ciliated, fibroblasts and myofibroblasts were significantly increased in IPF. We also found pDCs, ionocytes, and PNECs were significantly increased in IPF, although in very small proportions. Basal, club, goblet, aberrant basaloid cells, mature DCs, cDC1, B cells, regulatory T cells, mast cells, and ILCs were estimated to be zero (Fig. 3b).

### 3.3 Cell type proportion PCs as covariates

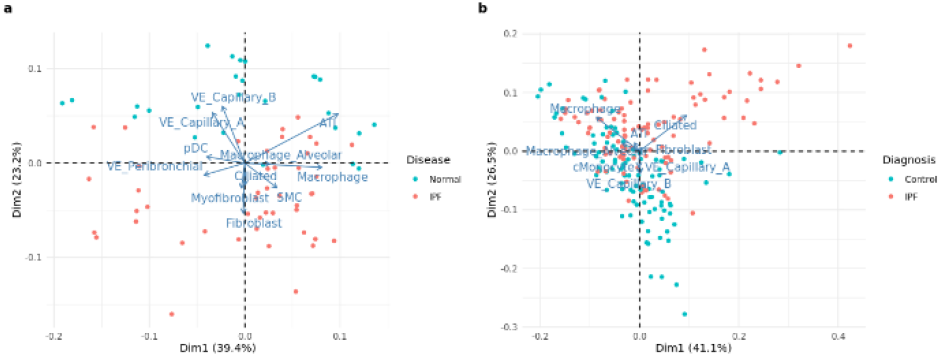

We then explored correcting the expression data for the varying cell type proportions using the top principal components of the proportions (Fig. 4, Fig. S3-S4). Correcting for the cell type proportions meant removing the effects of these varying proportions and the resulting differentially expressed genes are more likely to be associated with the disease. We considered doing two separate cell type proportions correction for the differential expression analysis: (1) using only PC1, and (2) using PC1-PC4.

There was an increase in the number of differentially expressed genes, 463 more genes after correcting for PC1 compared to without correction in the first dataset, and a slight decrease, 82 less genes, after correcting for PC1-PC4 compared to without correction (Fig. 2a, Fig. S5-S6). Meanwhile, in the second dataset, there are 426 less genes after PC1 correction compared to without correction, and 3380 less genes after PC1-PC4 correction compared to without correction (Fig. 2a, Fig. S7-S8).

Next, we performed pathway analysis after correction. Correcting for PC1 resulted in very similar results without correction, while correcting for PC1-PC4 resulted in few similar results without correction and with PC1 correction. For the first dataset, TGF-β regulation of extracellular matrix was the common pathway (Fig. 5a, 5b). For the second dataset, TGF-β regulation of extracellular matrix and axon guidance were the common pathways (Fig. 5c, 5d).

**Figure 5.**
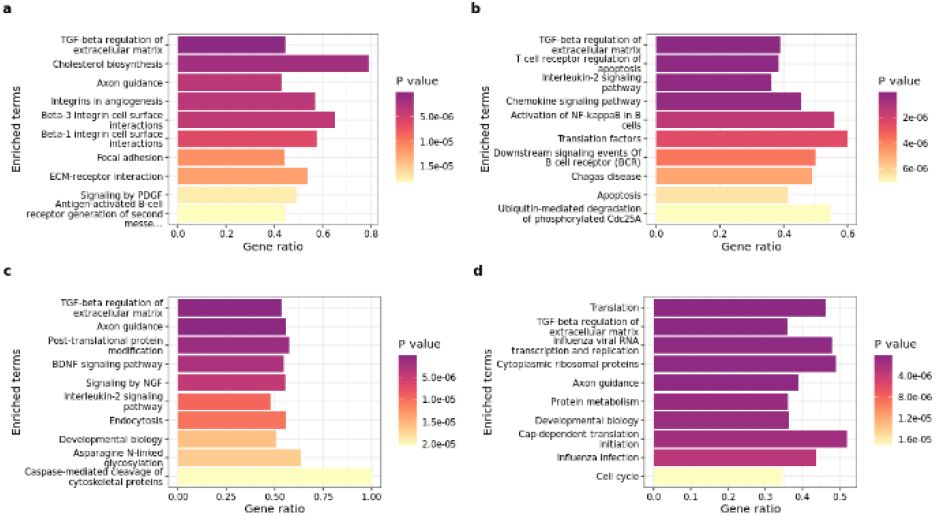
Pathway analysis for (a) GSE134692 with PC1 correction, (b) with PC1-PC4 correction, (c) GSE150910 with PC1 correction, (d) with PC1-PC4 correction.

### 3.4 Cell type-specific pathway analysis

We also explored cell type-specific pathway analysis for each cell type after correction (Fig. S9-S12). In the first dataset, fibroblasts pathways include extracellular matrix organization, collagen biosynthesis, syndecan 1 pathway, and beta-1 integrin cell surface interactions (Fig. 6b). In ATII cells, we identified interferon, EGF receptor (ErbB1) and insulin signaling pathways (Fig. 6c), while in VE capillary cells, pathways involving interleukin-4, muscarinic acetylcholine receptors and adrenoceptors were identified (Fig. 6d).

**Figure 6.**
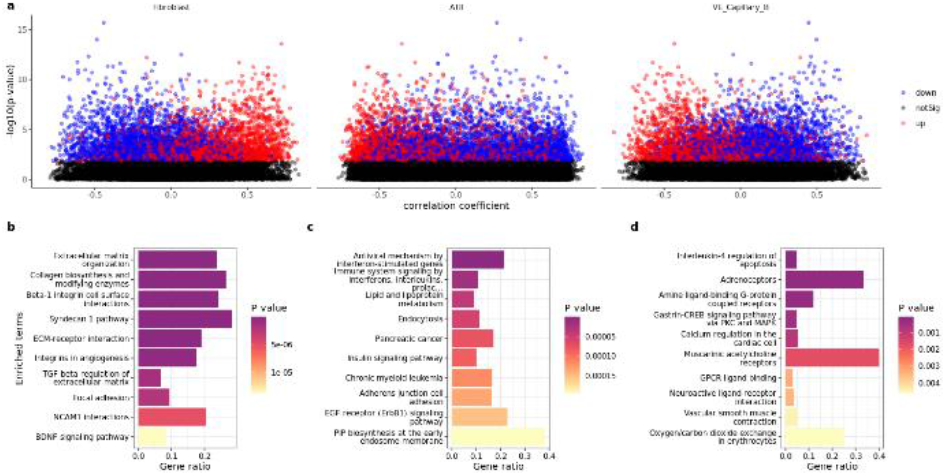
Cell type-specific pathway analysis for GSE134692 with PC1-PC4 correction. (a) log10p-value vs correlation plots for fibroblasts, ATII, VE capillary cells, (b) pathways for fibroblasts, (c) ATII cells, (d) VE capillary cells.

In the second dataset, fibroblasts pathways include extracellular matrix organization, syndecan 1, beta-1 integrin cell surface interactions, and collagen biosynthesis (Fig. 7b). In VE capillary cells, we identified pathways including endocytosis, agrin in postsynaptic differentiation, and calcium signaling (Fig. 7c), while in classical monocytes, pathways identified include RIG-I-like receptors signaling, TNFR2 signaling, NF-kB activation, interleukin-2 signaling (Fig. 7d).

**Figure 7.**
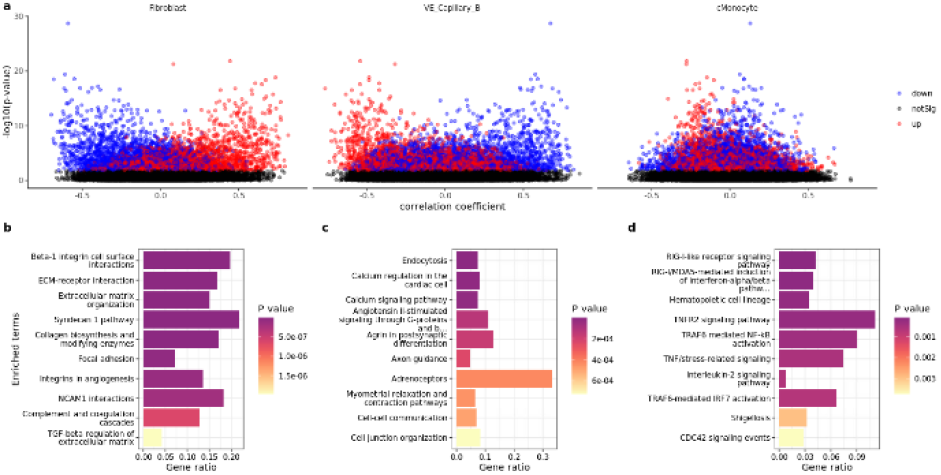
Cell type-specific pathway analysis for GSE150910 with PC1-PC4 correction. (a) log10p-value vs correlation plots for fibroblasts, VE capillary cells, classical monocytes, (b) pathways for fibroblasts, (c) VE capillary cells, (d) classical monocytes.

### 3.5 Comparison to cell type-specific pathways from scRNA-seq

We compared the cell type-specific pathways derived from deconvolution with those derived from scRNA-seq (Fig. 8, Fig. S13). A new IPF dataset was used (Habermann et al. 2020), independent of the one used with MuSiC. In lieu of ground truth, we chose six major pathways, previously established to be involved in IPF. In general, p-values were lower for these pathways using the deconvolution method.

**Figure 8.**
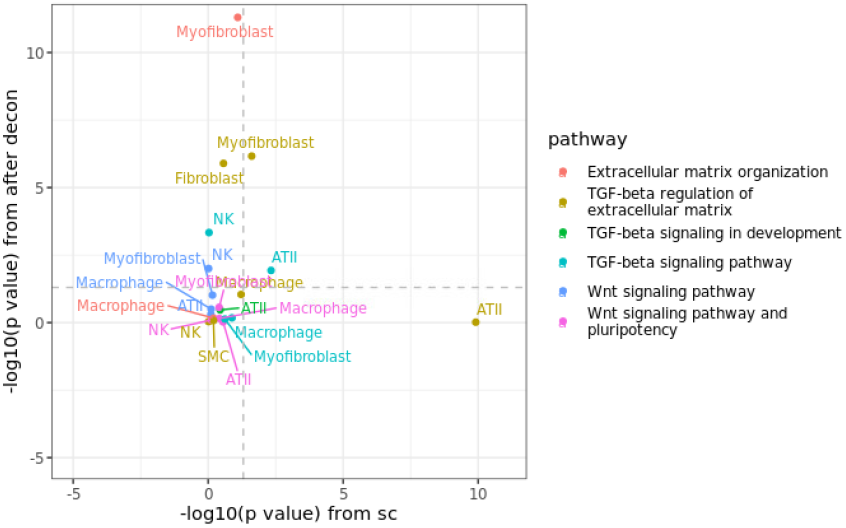
Validation of cell type-specific pathways in bulk data GSE134692.

The two approaches were concordant on *TGF-*β *regulation of extracellular matrix* in myofibroblasts and *TGF-*β *signaling* in ATII cells. Discrepancies between the methods include *TGF-*β *signaling/development* in NK cells, *TGF-*β *regulation of extracellular matrix* in fibroblasts, and *extracellular matrix organization* in myofibroblasts which were only detected using the deconvolution method. Conversely, the role of *TGF-*β *regulation of extracellular matrix* in ATII cells was only reported using the scRNA-seq method.

## 4 Discussion

### 4.1 Differences in cell-type proportions in IPF vs control samples

We performed deconvolution on IPF bulk RNA-seq data (Sivakumar et al. 2019; Furusawa et al. 2020) using MuSiC to estimate the cell type proportions. Since scRNA-seq data on IPF is readily available, using a reference-based deconvolution method such as MuSiC is relatively easy. MuSiC effectively uses the scRNA-seq data as reference for bulk RNA-seq data having different individuals (Wang et al. 2019). The method estimated most of the cell types and produced mostly consistent results across the two datasets. Estimated decreased proportions of cell types such as ATI and ATII epithelial cells, and VE capillary cells and increased proportions of cell types such as basal and goblet cells (although in very small proportions less than 10%), and ciliated cells are consistent with the study by Adams et al. 2020. However, MuSiC failed to estimate the proportions of some cell types and estimated them to be zero. In the second dataset (Furusawa et al. 2020), proportions of ATII cells, basal, club, and goblet cells were zero.

### 4.2 Disentangling gene regulation and changes in cell type proportions

Since bulk RNA-seq data contains a mixture of cell types and only represents the averaged expression of each gene across the cell types, the identification of differentially expressed genes is confounded by these varying cell type proportions (Cobos et al. 2018). Several studies had shown the effect and the importance of correcting for these cell types variation in the analysis of brain tissue (Patrick et al. 2020), using microarray data from mammary gland (Wang et al. 2006), using human blood and mouse kidney samples (Kong et al. 2019).

We performed PCA on the cell type proportions to reduce the number of variables for correction in the differential expression analysis. Reducing the number of covariates by PCA is a more ideal approach since there are over 20 cell types estimated by deconvolution and the cell type proportions are correlated with each other (Kong et al. 2019). We considered using PC1 for correction since this always explains the most variation, and also PC1-PC4, for comparison.

### 4.3 Cell type-specific pathway analysis

We explored the potential of further using the deconvolution results by performing cell type-specific pathway analysis. Correcting for only one PC yielded similar pathways to the original pathway analysis. Correcting for PC1-PC4 brought a series of new pathways to the surface. To assess the validity of the proposed pathways, we performed a literature analysis of these pathways in the context of IPF.

We identified the EGF receptor (ErbB1) signaling pathway as being modified in IPF and related to ATII cells. This is consistent with Tzouvelekis et al. 2013, where they investigated the role of EGFR in lung fibrosis. ErbB1 is upregulated in lung epithelial cells which is associated with fibrosis development (Tzouvelekis et al. 2013; Schramm, Schaefer, and Wygrecka 2022). We also identified the TGF-β regulation to be involved in IPF and related to fibroblasts and myofibroblasts (Fig. S10-S13). This pathway is also involved in the development of fibrosis and is also known to be related to other cell types such as macrophages, endothelial cells, and epithelial cells (Fernandez and Eickelberg 2012; Frangogiannis 2020). Moreover, our method found TGF-β signaling to be active in endothelial cells (Fig. S10, S12). The activation of TGF-β signaling in endothelial cells was studied in mice (Wermuth et al. 2017), where they found that upon activation, it caused fibrosis not only in the lungs but also skin, liver, kidney, and heart, and they described it to be similar with the fibrotic characteristics of systemic sclerosis.

We also identified neurotrophin signaling pathway as modified in IPF and related to ATII cells (Fig. S10). Neurotrophins are molecules involved in the nervous system development, but they are also involved and expressed in the lungs. They include nerve growth factor (NGF), brain-derived neurotrophic factor (BDNF), neurotrophin-3 (NT-3), and neurotrophin-4 (NT-4), and are expressed in lung cells such as smooth muscle cells, fibroblasts and vascular endothelial cells, and immune cells (Hoyle 2003; Prakash et al. 2010; Rubin et al. 2021). Neurotrophins are known to be involved in certain lung functions occurring at different cell types, such as development, airway remodeling, angiogenesis, cell migration and proliferation (Hoyle 2003; Prakash et al. 2010; Rubin et al. 2021). Moreover, the expression of NT-4/5 and its receptor TrkB were increased in ATII cells in the lungs of IPF patients, and this was also evident in the lungs of mice where pulmonary fibrosis was induced (Avcuoglu et al., 2011).

Wnt signaling was also identified to be modified in IPF and related to VE capillary cells (Fig. S10). This pathway is involved in vascular development and angiogenesis in endothelial cells, where different Wnt components have different specific biological functions during development, such as cell proliferation, cell differentiation, vessel assembly in endothelial cells (Franco, Liebner, and Gerhardt 2009). In the lungs, in addition to development, it plays a major role in repair and regeneration, and disease progression (Aros, Pantoja, and Gomperts 2021). Another study showed that Wnt and TGF pathways were activated by the overexpression of SREBP2 in endothelial cells in IPF (Martin et al. 2021). The overexpression also caused phenotypic and epigenetic change of endothelial cells, which then led to increased ECM deposition, proliferation, and exacerbation of pulmonary fibrosis (Martin et al. 2021). We compared the deconvolution-derived cell type-specific pathways with those generated directly in scRNA-seq. This point of comparison is challenging because of lack of ground truth. Given the lack of depth of sequencing in scRNA-seq, it would be premature to consider it a gold standard for this comparison. Given that the deconvolution-based approach delivered lower p-values for many pathway types, this could be an indication of greater sensitivity, consistent with the greater depth of sequencing in the bulk RNA-seq. However, there is no objective way to rule out false positives from the deconvolution method, and therefore we rely on the literature based validation discussed above to conclude that cell type-specific pathway analysis based on deconvolution has potential but requires validation from independent, targeted experiments.

We have shown a new analysis pipeline for bulk RNA-seq data involving deconvolution, which allowed us to estimate cell type proportions and to further perform cell type-specific pathway analysis. The proposed pipeline efficiently enables the re-analysis of several bulk datasets, which opens possibilities to gain new knowledge about a specific disease that will otherwise be missed.

## References

[dataset]* Adams TS, Schupp, JC, Poli, S et al. Single-cell RNA-seq reveals ectopic and aberrant lung-resident cell populations in idiopathic pulmonary fibrosis. Sci Adv. 2020, 6(28): eaba1983. 10.1126/sciadv.aba1983.

Aros, CJ, Pantoja, CJ, and Gomperts, BN. Wnt signaling in lung development, regeneration, and disease progression. Commun Biol 2021, 4(601), 10.1038/s42003-021-02118-w.

Avcuoglu, S, Wygrecka, M, Marsh, LM et al. Neurotrophic tyrosine kinase receptor B/ neurotrophin 4 signaling axis is perturbed in clinical and experimental pulmonary fibrosis. Am. J. Respir. Cell Mol. Biol. 2011, 45: 768–780, 10.1165/rcmb.2010-0195OC.

Chen, EY, Tan, CM, Kou, Y et al. Enrichr: interactive and collaborative HTML5 gene list enrichment analysis tool. BMC Bioinformatics, 2013: 14(128). 10.1186/1471-2105-14-128.

Cobos, F, Vandesompele, J, Mestdagh, P et al. Computational deconvolution of transcriptomics data from mixed cell populations. Bioinformatics 2018, 34(11): 1969–1979. 10.1093/bioinformatics/bty019.

Fernandez, IE and Eickelberg, O. The Impact of TGF-β on Lung Fibrosis. Proceedings of the American Thoracic Society 2012, 9(3), 111–116. 10.1513/pats.201203-023aw.

Franco, CA, Liebner, S, and Gerhardt, H. Vascular morphogenesis: a Wnt for every vessel? Current Opinion in Genetics & Development 2009, 19(5): 476–483. 10.1016/j.gde.2009.09.004.

Frangogiannis, NG. Transforming growth factor–β in tissue fibrosis. J Exp Med 2020, 217 (3): e20190103. 10.1084/jem.20190103.

[dataset]* Furusawa, H, Cardwell, JH, Okamoto, T et al. Chronic Hypersensitivity Pneumonitis, an Interstitial Lung Disease with Distinct Molecular Signatures. Am. J. Respir. Crit. Care Med. 2020, 202: 1430–1444. 10.1164/rccm.202001-0134OC.

Ghosh, AJ, Hobbs, BD, Yun, JH et al. Lung tissue shows divergent gene expression between chronic obstructive pulmonary disease and idiopathic pulmonary fibrosis. Respir Res 2022, 23, 97. 10.1186/s12931-022-02013-w.

[dataset]* Habermann, AC, Gutierrez, AJ, Bui, LT et al. Single-cell RNA sequencing reveals profibrotic roles of distinct epithelial and mesenchymal lineages in pulmonary fibrosis. Sci Adv. 2020, 6(28): eaba1972. 10.1126/sciadv.aba1972.

Hao, Y, Hao, S, Andersen-Nissen, E et al. Integrated analysis of multimodal single-cell data. Cell 2021, 10.1016/j.cell.2021.04.048.

Hoyle, GW. Neurotrophins and lung disease. Cytokine & Growth Factor Reviews 2003, 14(6): 551–558. 10.1016/s1359-6101(03)00061-3.

Huang, Y, Guzy, R, Ma, S-F et al. Central lung gene expression associates with myofibroblast features in idiopathic pulmonary fibrosis. BMJ Open Resp Res 2023, 10: e001391. 10.1136/bmjresp-2022-001391.

Jew, B, Alvarez, M, Rahmani, E et al. Accurate estimation of cell composition in bulk expression through robust integration of single-cell information. Nat Commun 2020, 11: 1971. 10.1038/s41467-020-15816-6.

Kanter, I and Kalisky, T. Single cell transcriptomics: methods and applications. Front Oncol. 2015, 5: 53. 10.3389/fonc.2015.00053.

Kong, Y, Rastogi, D, Seoighe, C et al. Insights from deconvolution of cell subtype proportions enhance the interpretation of functional genomic data. PLoS ONE 2019, 14(4): e0215987. 10.1371/journal.pone.0215987.

Martin, M, Zhang, J, Miao, Y et al. Role of endothelial cells in pulmonary fibrosis via SREBP2 activation. JCI Insight 2021, 6(22): e125635. 10.1172/jci.insight.125635.

Menden, K, Marouf, M, Oller, S et al. Deep learning-based cell composition analysis from tissue expression profiles. Sci Adv. 2020, 6(30): eaba2619. 10.1126/sciadv.aba2619.

Newman, AM, Steen, CB, Liu, CL et al. Determining cell type abundance and expression from bulk tissues with digital cytometry. Nat Biotechnol 2019, 37: 773–782. 10.1038/s41587-019-0114-2.

Patrick, E, Taga, M, Ergun, A et al. Deconvolving the contributions of cell-type heterogeneity on cortical gene expression. PLoS Comput Biol 2020, 16(8): e1008120. 10.1371/journal.pcbi.1008120.

Pellegrino-Coppola, D, Claringbould, A, Stutvoet, M et al. Correction for both common and rare cell types in blood is important to identify genes that correlate with age. BMC Genomics 2021, 22(1): 184. 10.1186/s12864-020-07344-w.

Prakash, Y, Thompson, MA, Meuchel, L et al. Neurotrophins in lung health and disease. Expert Rev Respir Med. 2010, 4(3): 395–411. 10.1586/ers.10.29.

Reyfman, PA, Walter, JM, Joshi, N et al Single-Cell Transcriptomic Analysis of Human Lung Provides Insights into the Pathobiology of Pulmonary Fibrosis. American Journal of Respiratory and Critical Care Medicine 2018, 199. 10.1164/rccm.201712-2410oc.

Ritchie, ME, Phipson, B, Wu, D et al. limma powers differential expression analyses for RNA-sequencing and microarray studies. Nucleic Acids Research 2015, 43(7): e47. 10.1093/nar/gkv007.

Robinson, MD, McCarthy, DJ, and Smyth, GK. edgeR: a Bioconductor package for differential expression analysis of digital gene expression data. Bioinformatics 2010, 26(1), 139-140. 10.1093/bioinformatics/btp616.

Rubin, L, Stabler, CT, Schumacher-Klinger, A et al. Neurotrophic factors and their receptors in lung development and implications in lung diseases. Cytokine & Growth Factor Reviews 2021, 59: 84–94. 10.1016/j.cytogfr.2021.01.008.

Satija, R, Farrell, JA, Gennert, D et al. Spatial reconstruction of single-cell gene expression data. Nature Biotechnology 2015, 33: 495–502. 10.1038/nbt.3192.

[dataset]* Sivakumar, P, Thompson, JR, Ammar, R et al. RNA sequencing of transplant-stage idiopathic pulmonary fibrosis lung reveals unique pathway regulation. ERJ Open Res. 2019, 5(3):00117–2019. 10.1183/23120541.00117-2019.

Stegle, O, Teichmann, SA, and Marioni, JC. Computational and analytical challenges in single-cell transcriptomics. Nature Reviews Genetics 2015, 16(3), 133–145. 10.1038/nrg3833.

Torroja, C and Sanchez-Cabo, F. Digitaldlsorter: Deep-Learning on scRNA-Seq to Deconvolute Gene Expression Data. Frontiers in Genetics 2019, 10. 10.3389/fgene.2019.00978.

Tzouvelekis, A, Ntolios, P, Karameris, A et al. Increased expression of epidermal growth factor receptor (EGF-R) in patients with different forms of lung fibrosis. Biomed Res Int. 2013, 2013: 654354. 10.1155/2013/654354.

Wang, M, Master, SR, and Chodosh, LA. Computational expression deconvolution in a complex mammalian organ. BMC Bioinformatics 2006, 7: 328. 10.1186/1471-2105-7-328.

Wang, X, Park, J, Susztak, K et al. Bulk tissue cell type deconvolution with multi-subject single-cell expression reference. Nat Commun 2019, 10: 380. 10.1038/s41467-018-08023-x.

Wermuth, P, Carney, K, Mendoza, F et al. Endothelial cell-specific activation of transforming growth factor-β signaling in mice induces cutaneous, visceral, and microvascular fibrosis. Lab Invest 2017, 97(7): 806–818. 10.1038/labinvest.2017.23

