## Supplemental Figures for "Extending the capabilities of deconvolution to provide cell type specific pathway analysis of bulk RNA-seq data for idiopathic pulmonary fibrosis"

### Supplementary Figures

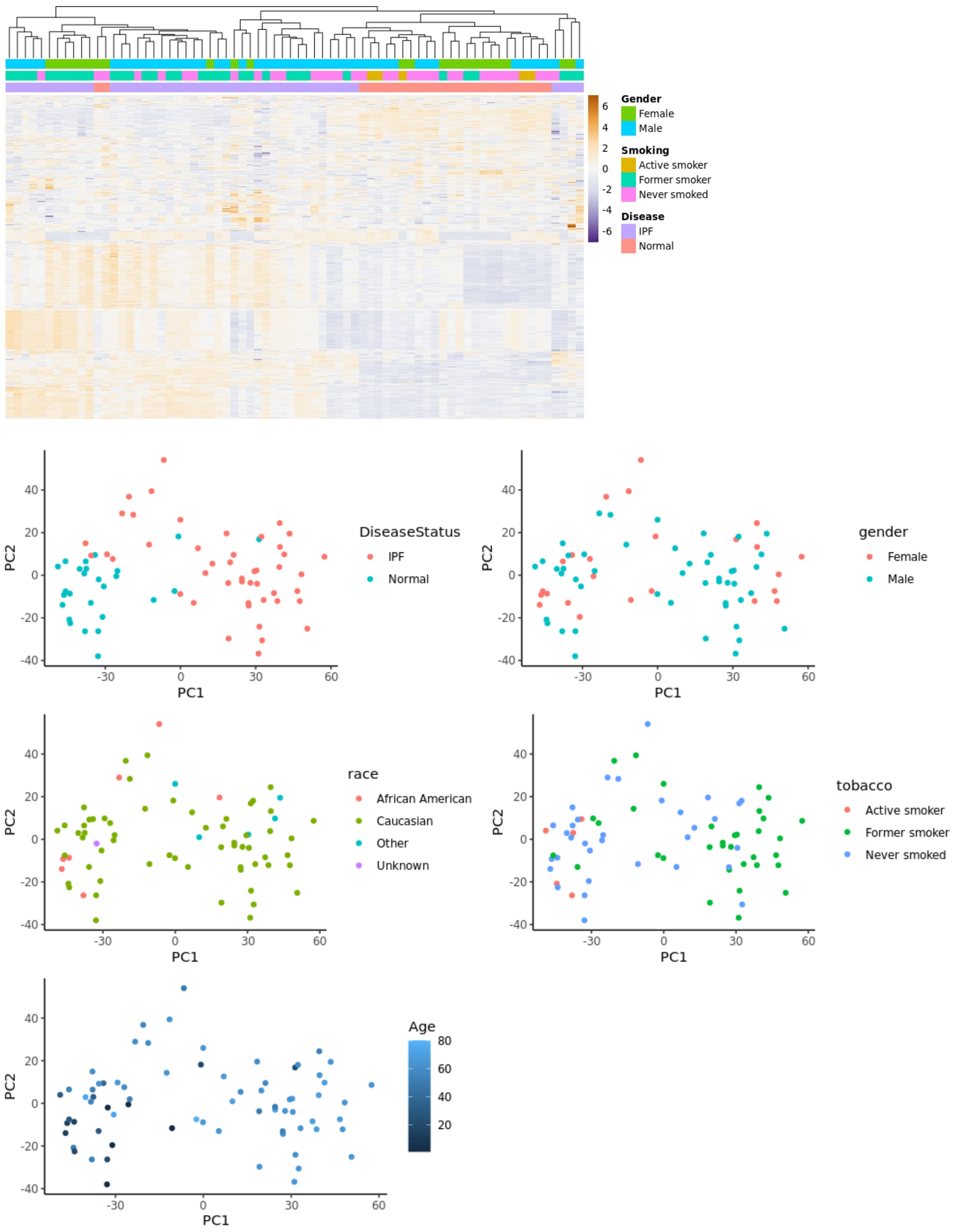


**Fig. S1.** Pre-processing and unsupervised analysis of GSE134692.

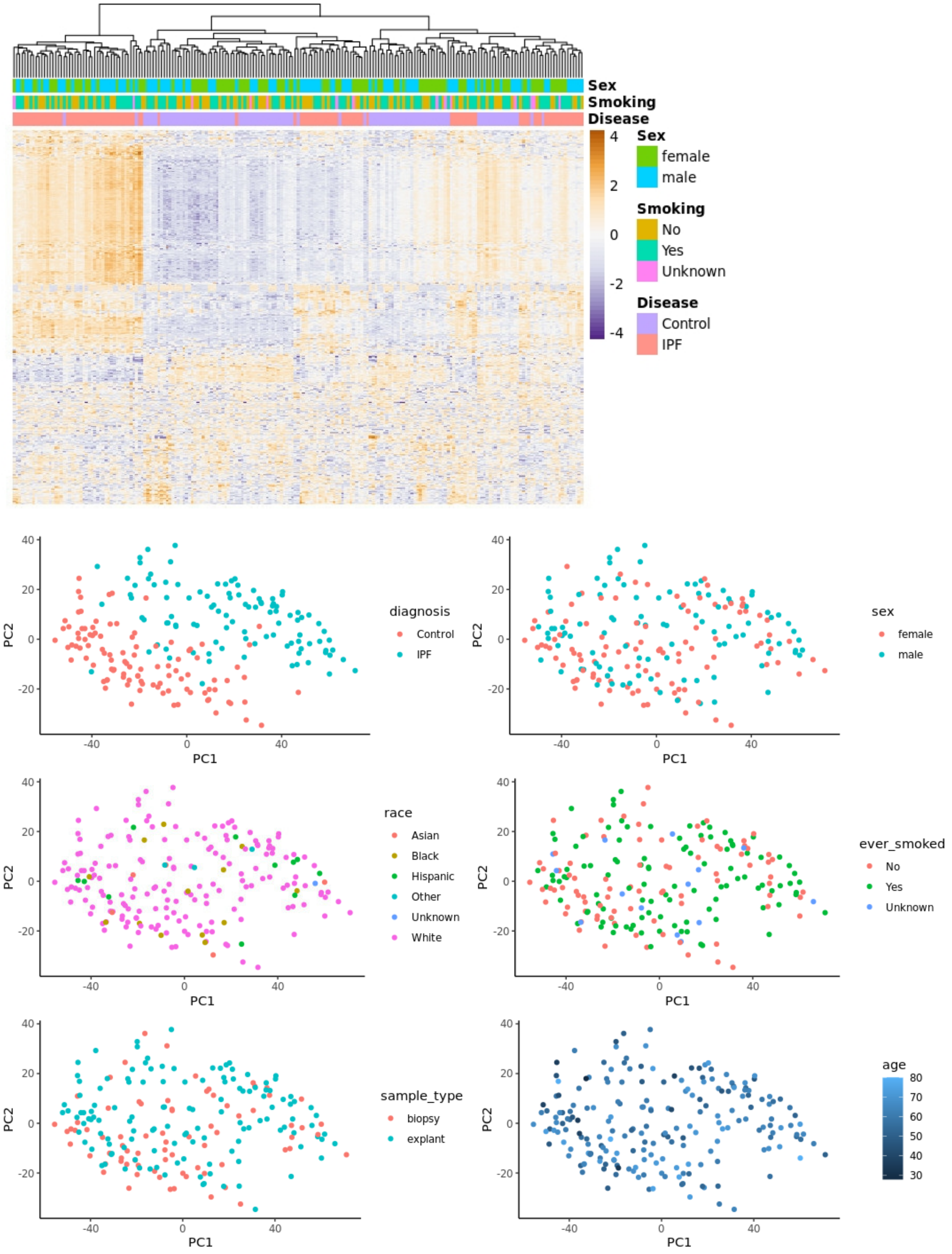


**Fig. S2.** Pre-processing and unsupervised analysis of GSE150910.

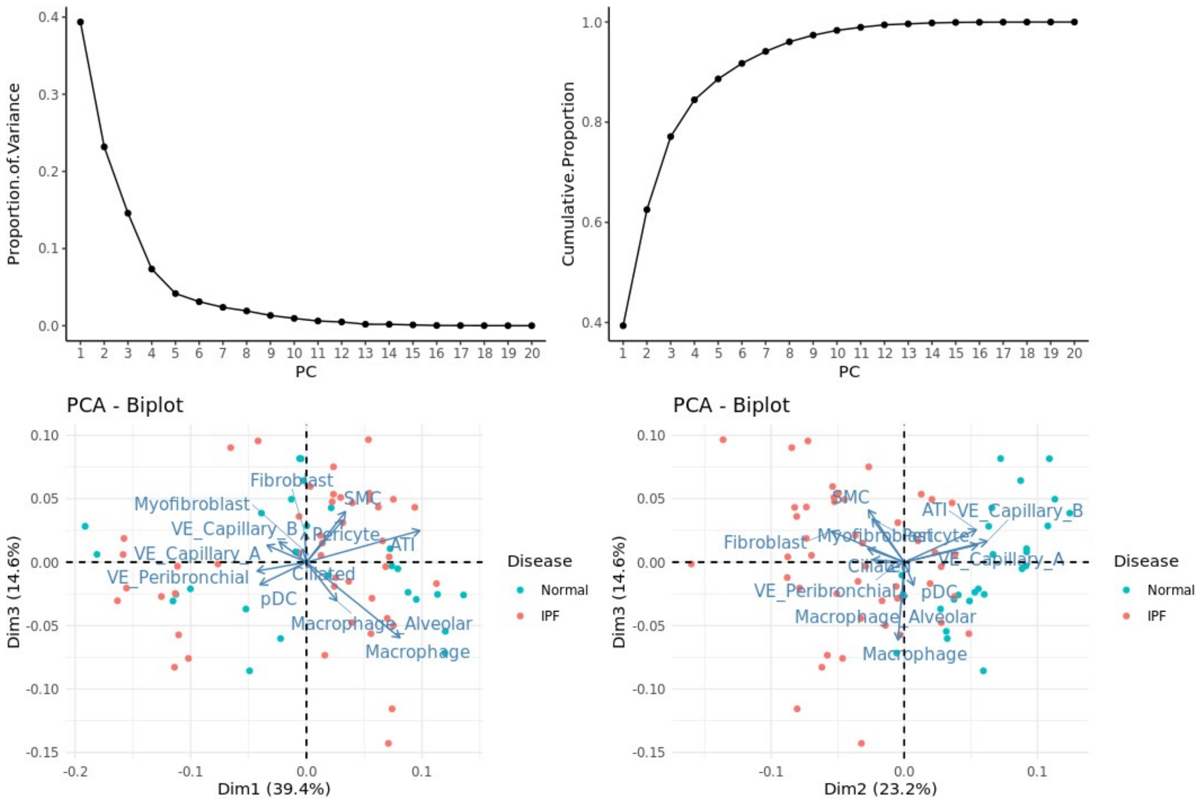


**Fig. S3.** PCA of the estimated cell type proportions in GSE134692. Cell types with zero proportions for all samples are not included in the PCA.

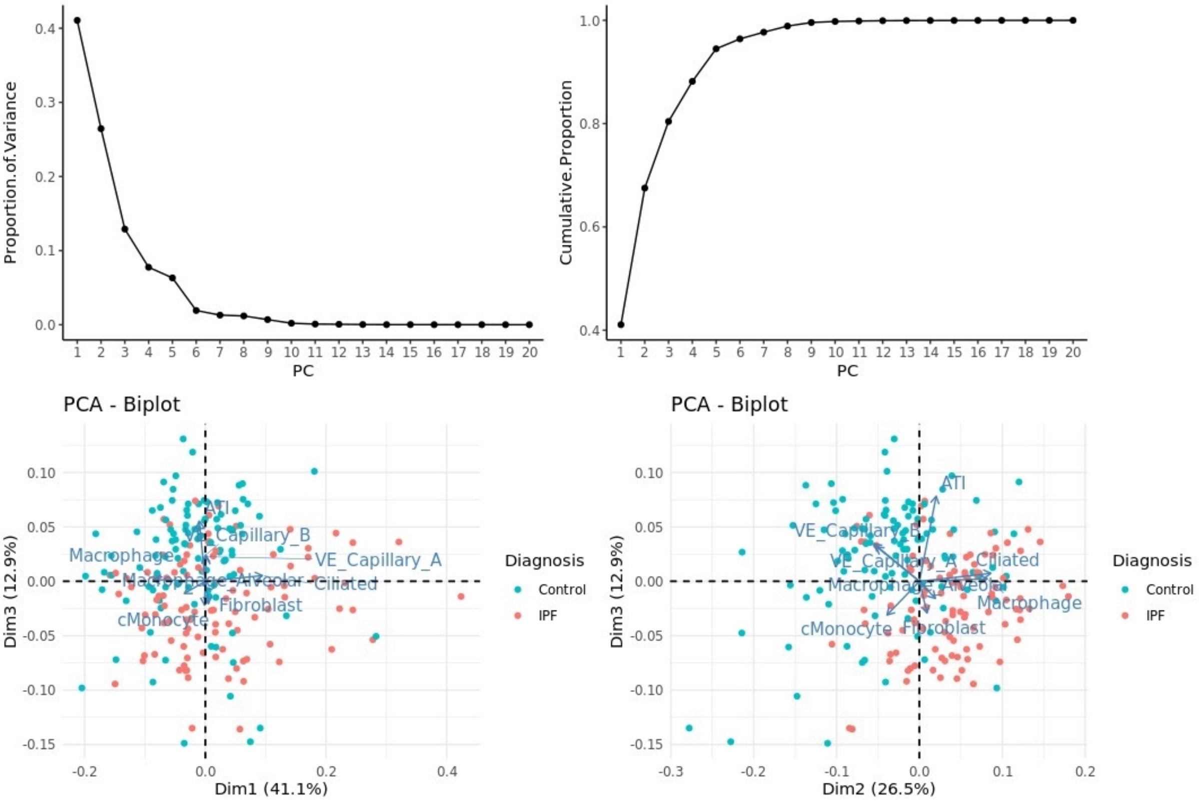


**Fig. S4.** PCA of the estimated cell type proportions in GSE150910. Cell types with zero proportions for all samples are not included in the PCA.

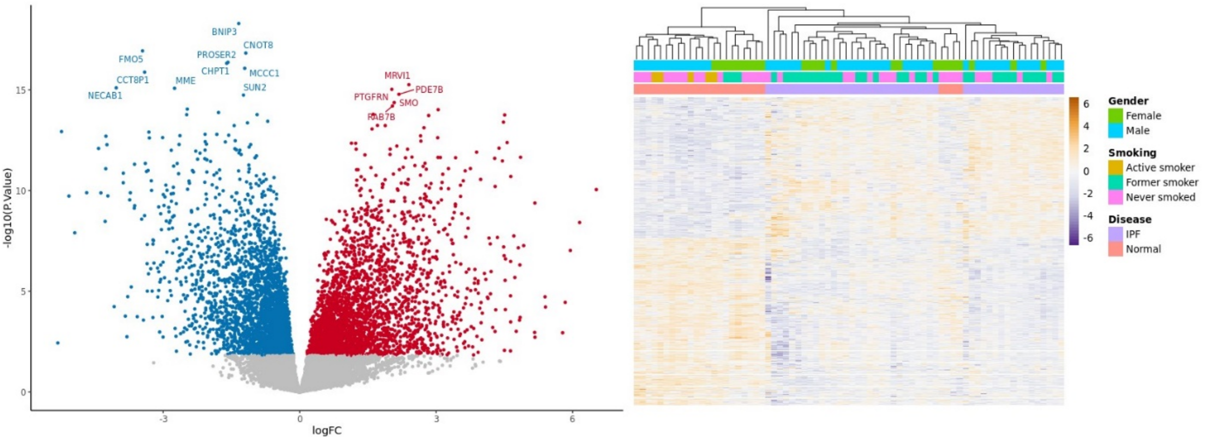


**Fig. S5.** Differential expression analysis results for GSE134692 after PC1 correction. Volcano plot and heatmap of DE genes.

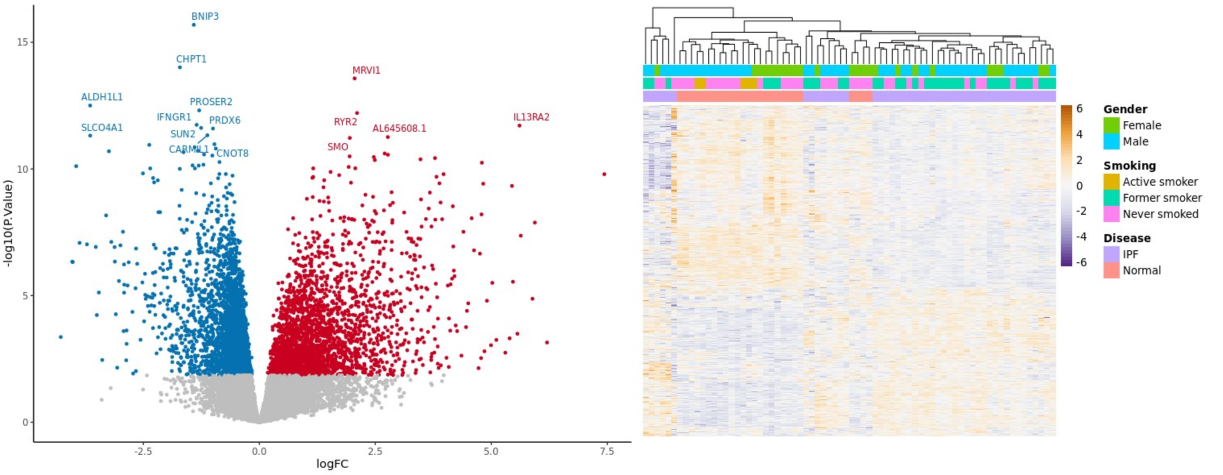


**Fig. S6.** Differential expression analysis results for GSE134692 after PC1-PC4 correction. Volcano plot and heatmap of DE genes.

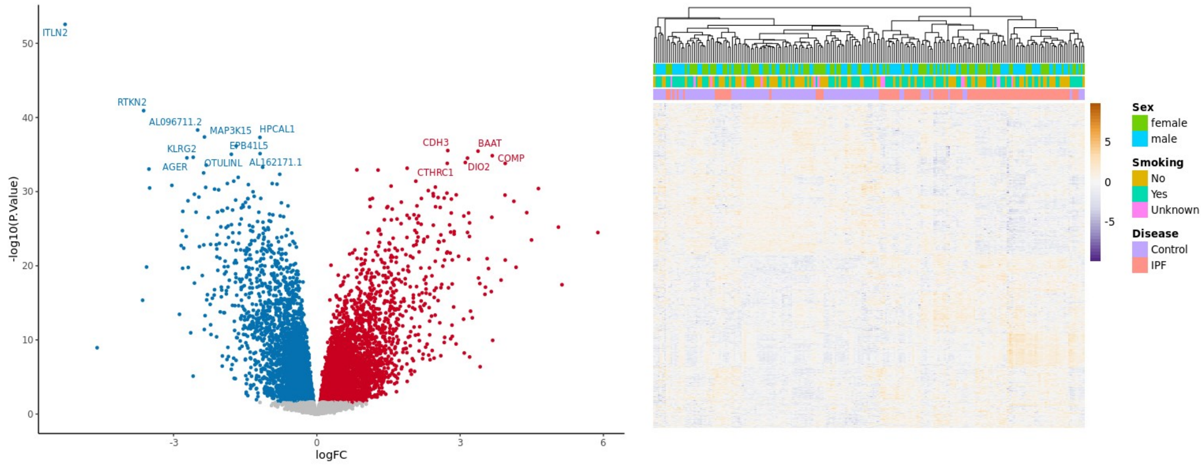


**Fig. S7.** Differential expression analysis results for GSE1150910 after PC1 correction. Volcano plot and heatmap of DE genes.

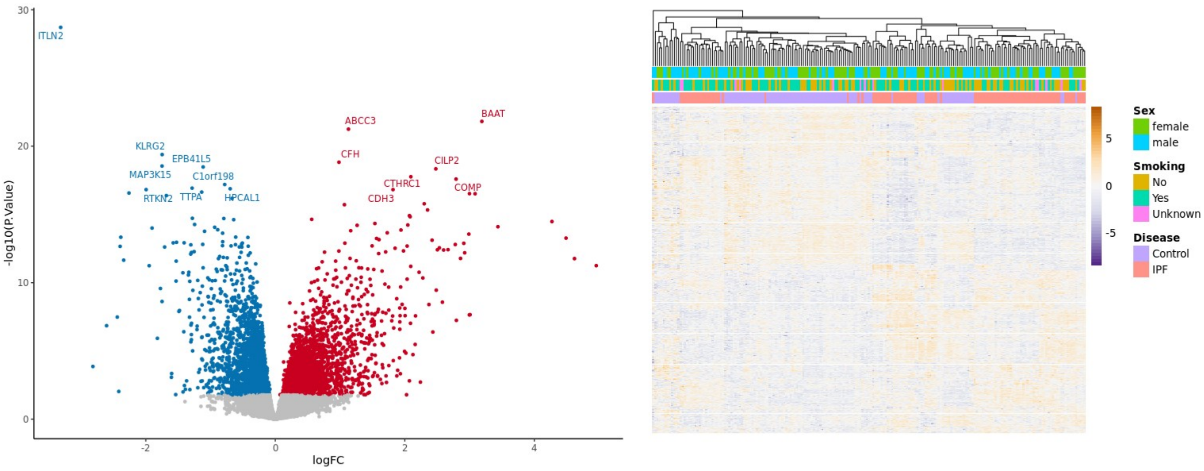


**Fig. S8.** Differential expression analysis results for GSE1150910 after PC1-PC4 correction. Volcano plot and heatmap of DE genes.

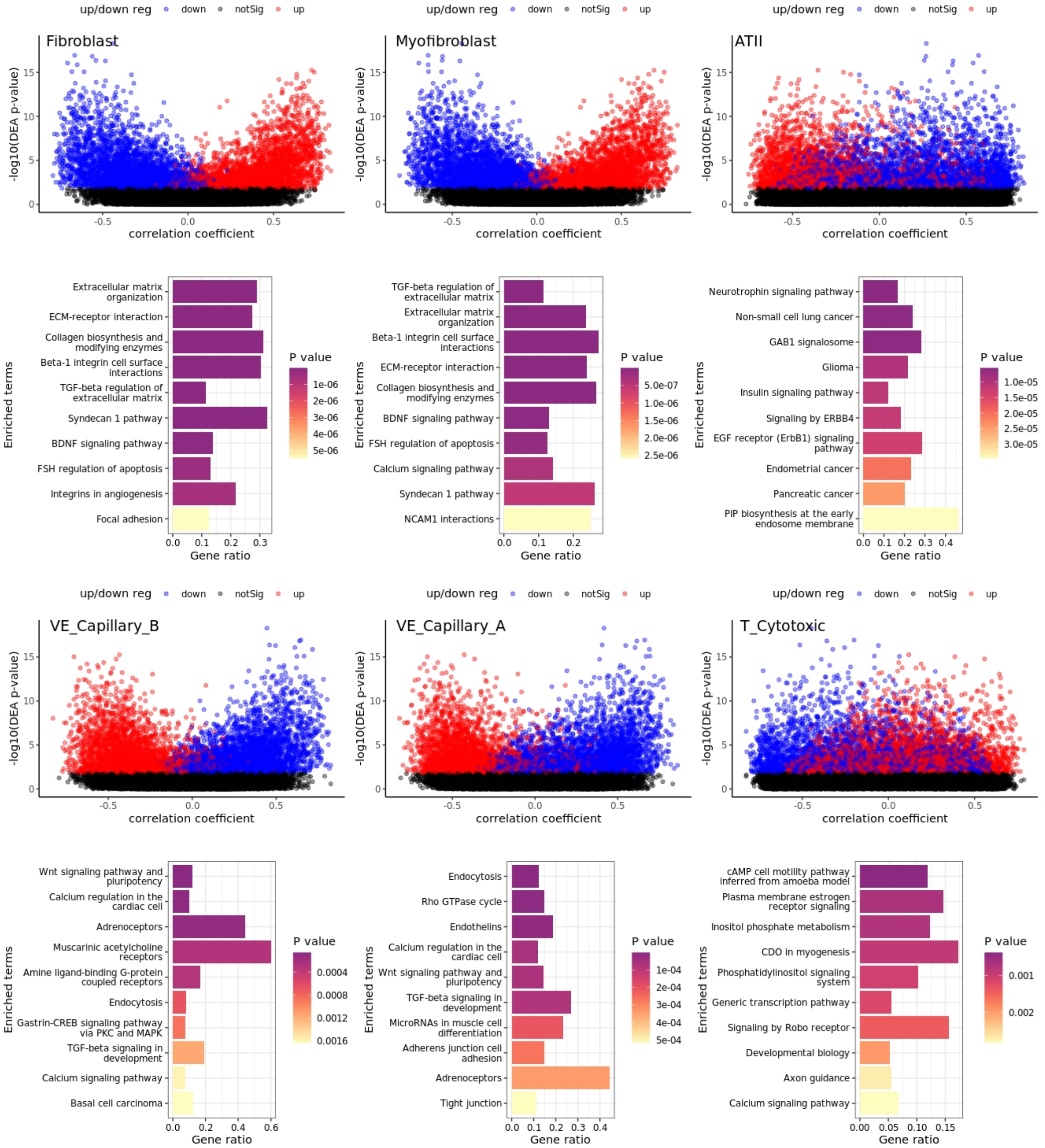


**Fig. S9.** Cell-type specific pathway analysis. $\log_{10}$p-value vs. correlation plots and pathways per cell type, after PC1 correction for GSE134692. Pathway result for each cell type is directly below its scatter plot.

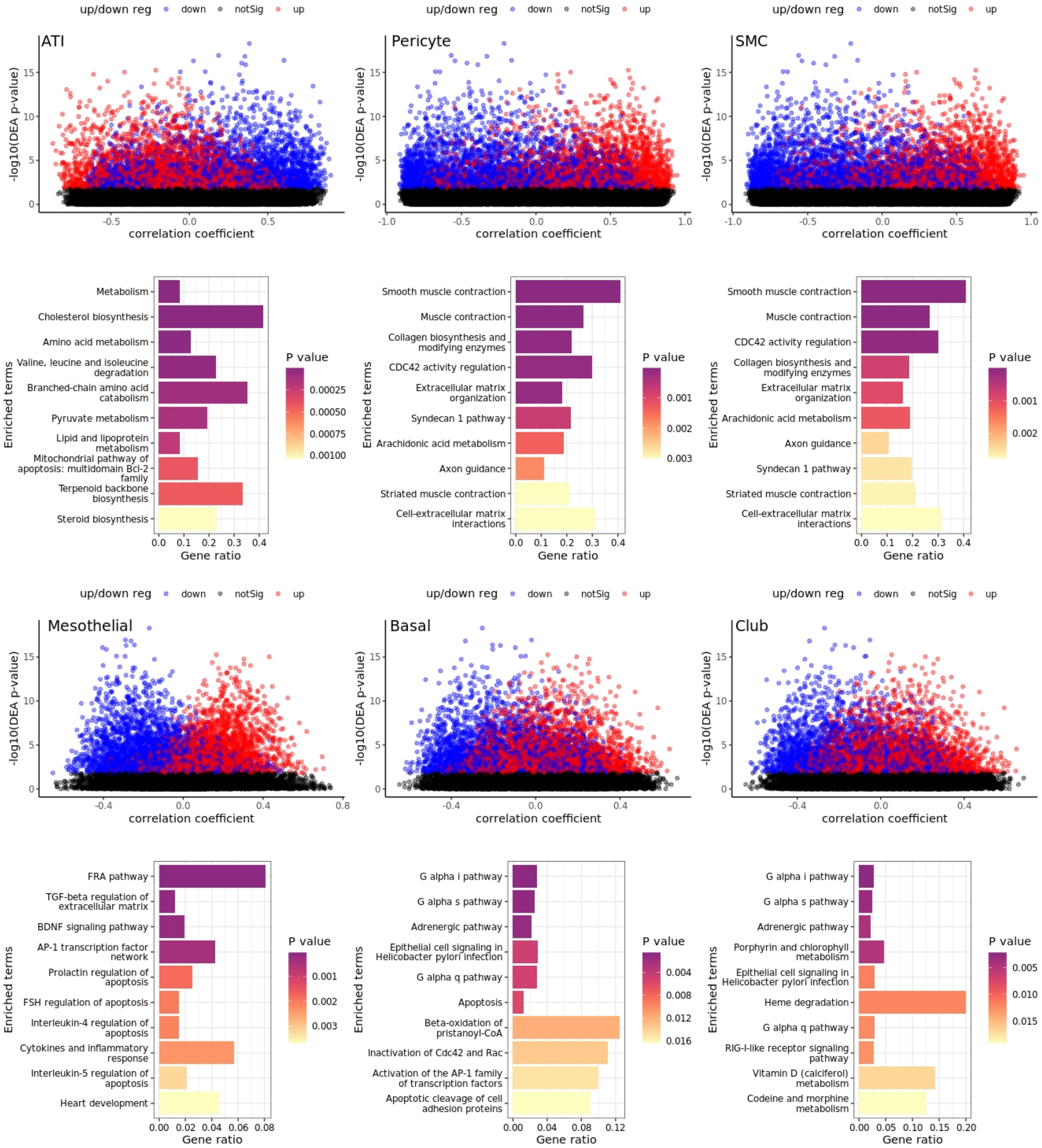


**Fig. S9 (cont'd).** Cell-type specific pathway analysis. $\log_{10}$p-value vs. correlation plots and pathways per cell type, after PC1 correction for GSE134692. Pathway result for each cell type is directly below its scatter plot.

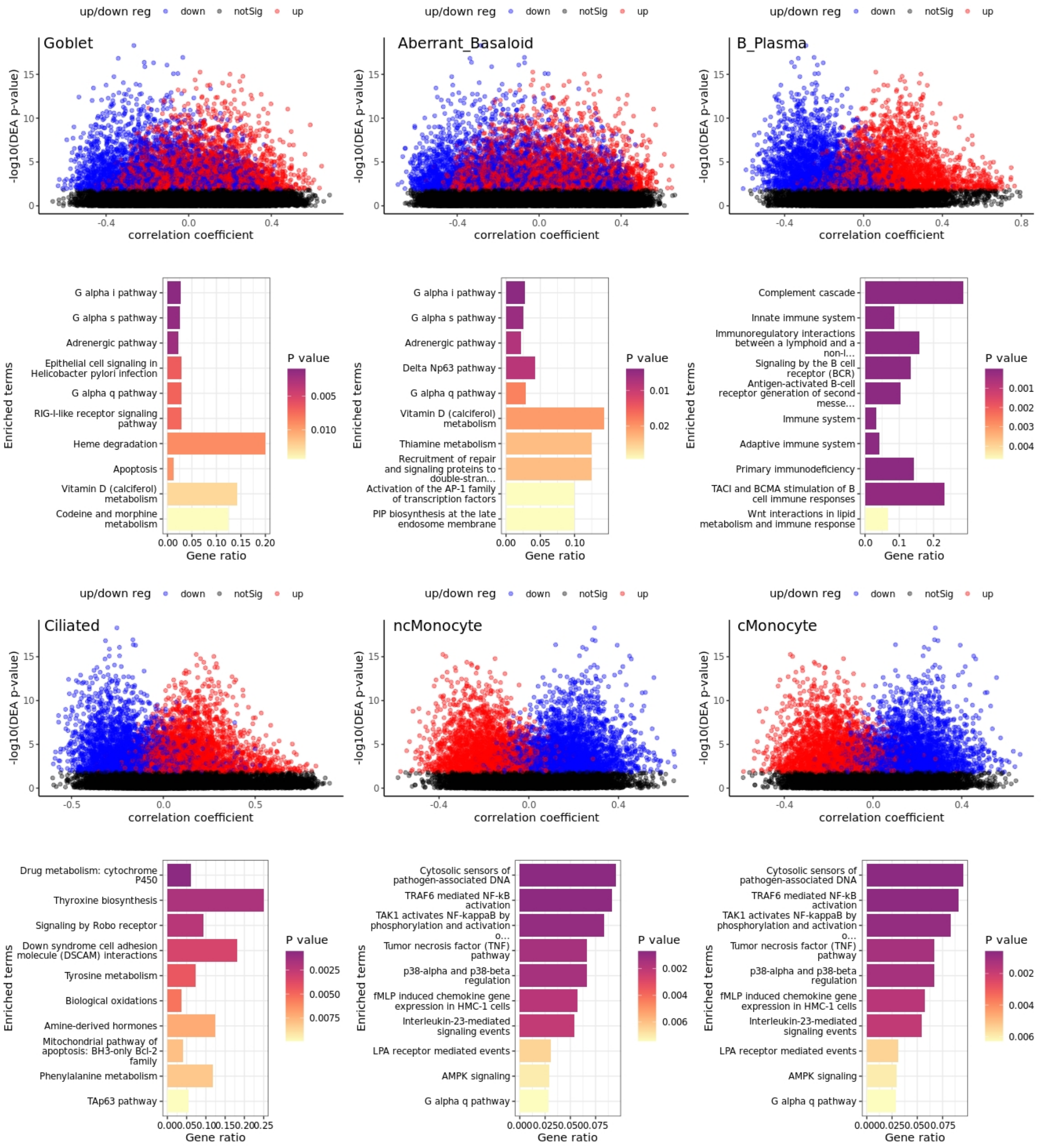


**Fig. S9 (cont'd).** Cell-type specific pathway analysis. $\log_{10}$p-value vs. correlation plots and pathways per cell type, after PC1 correction for GSE134692. Pathway result for each cell type is directly below its scatter plot.

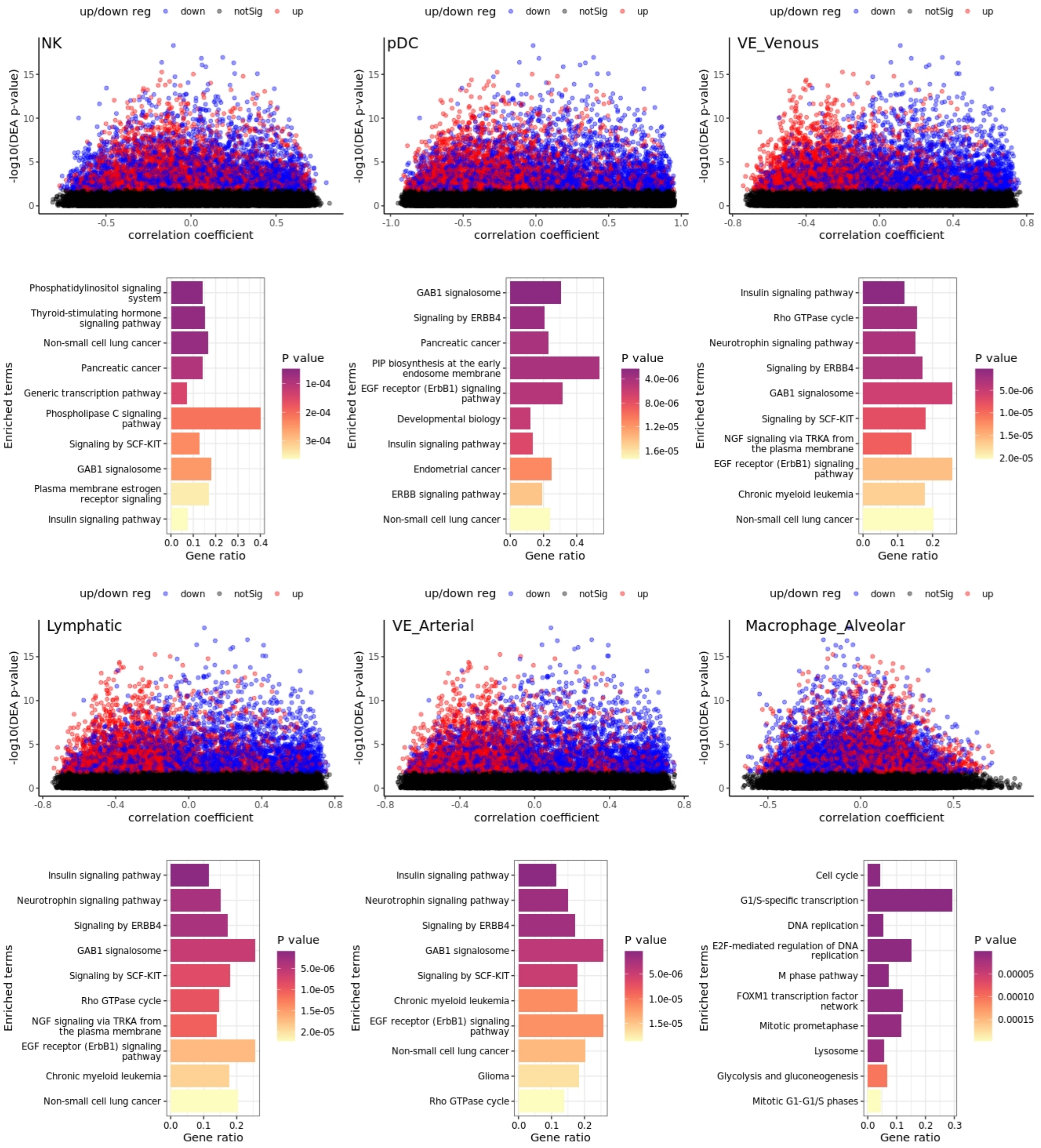


**Fig. S9 (cont'd).** Cell-type specific pathway analysis. $\log_{10}$p-value vs. correlation plots and pathways per cell type, after PC1 correction for GSE134692. Pathway result for each cell type is directly below its scatter plot.

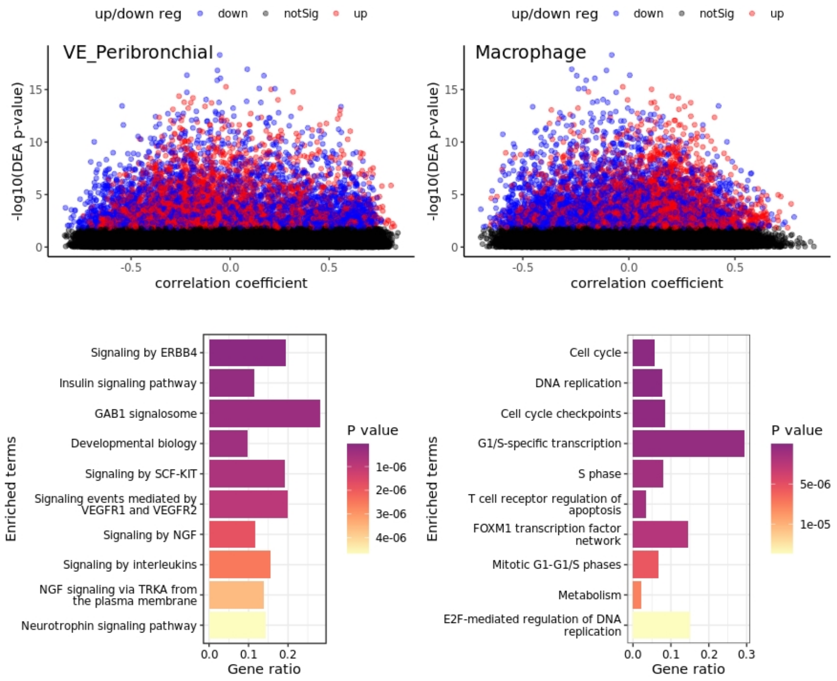


**Fig. S9 (cont'd).** Cell-type specific pathway analysis. $\log_{10}$p-value vs. correlation plots and pathways per cell type, after PC1 correction for GSE134692. Pathway result for each cell type is directly below its scatter plot.

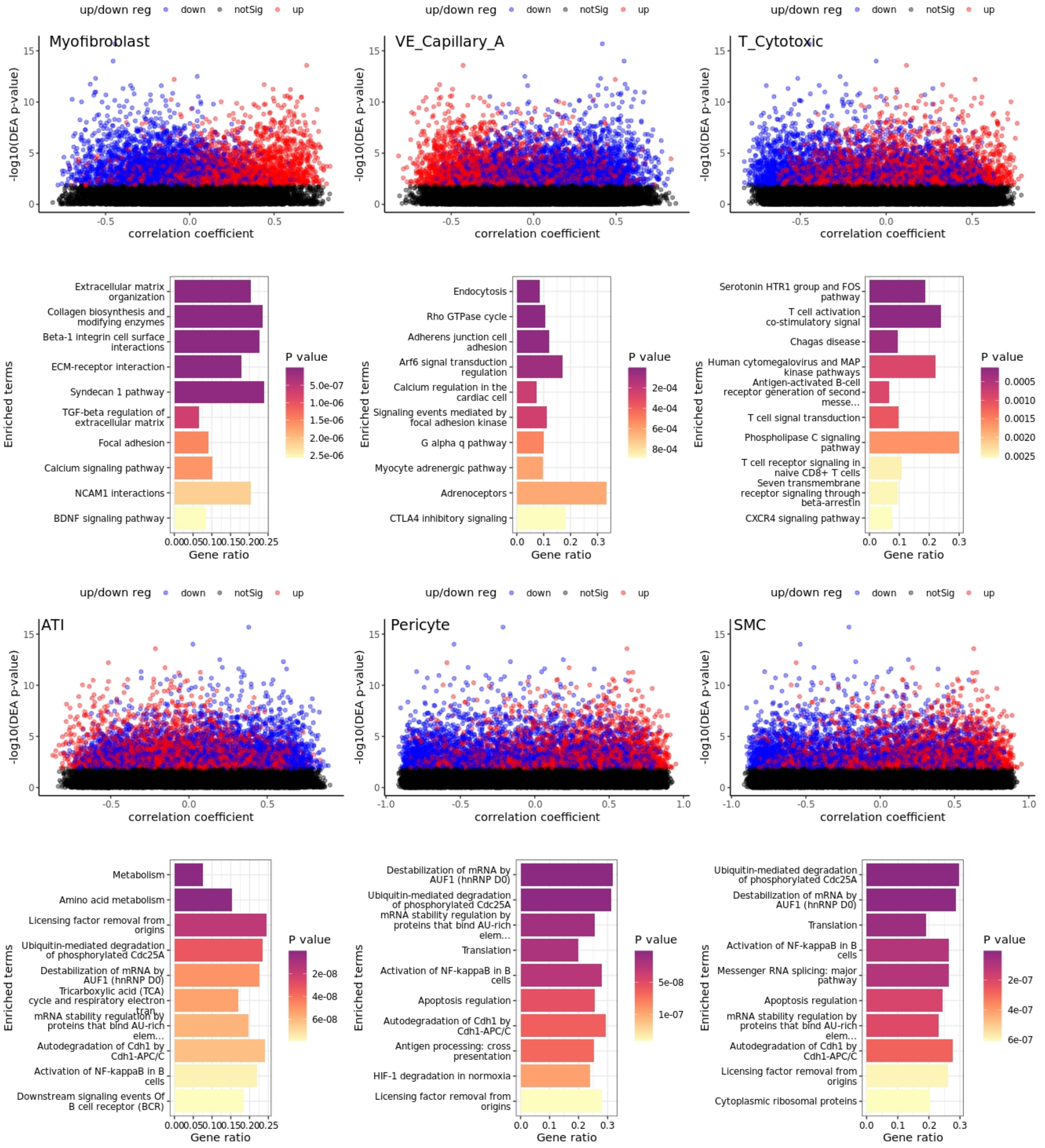


**Fig. S10.** Cell-type specific pathway analysis. $\log_{10}$p-value vs. correlation plots and pathways per cell type, after PC1-PC4 correction for GSE134692. Pathway result for each cell type is directly below its scatter plot.

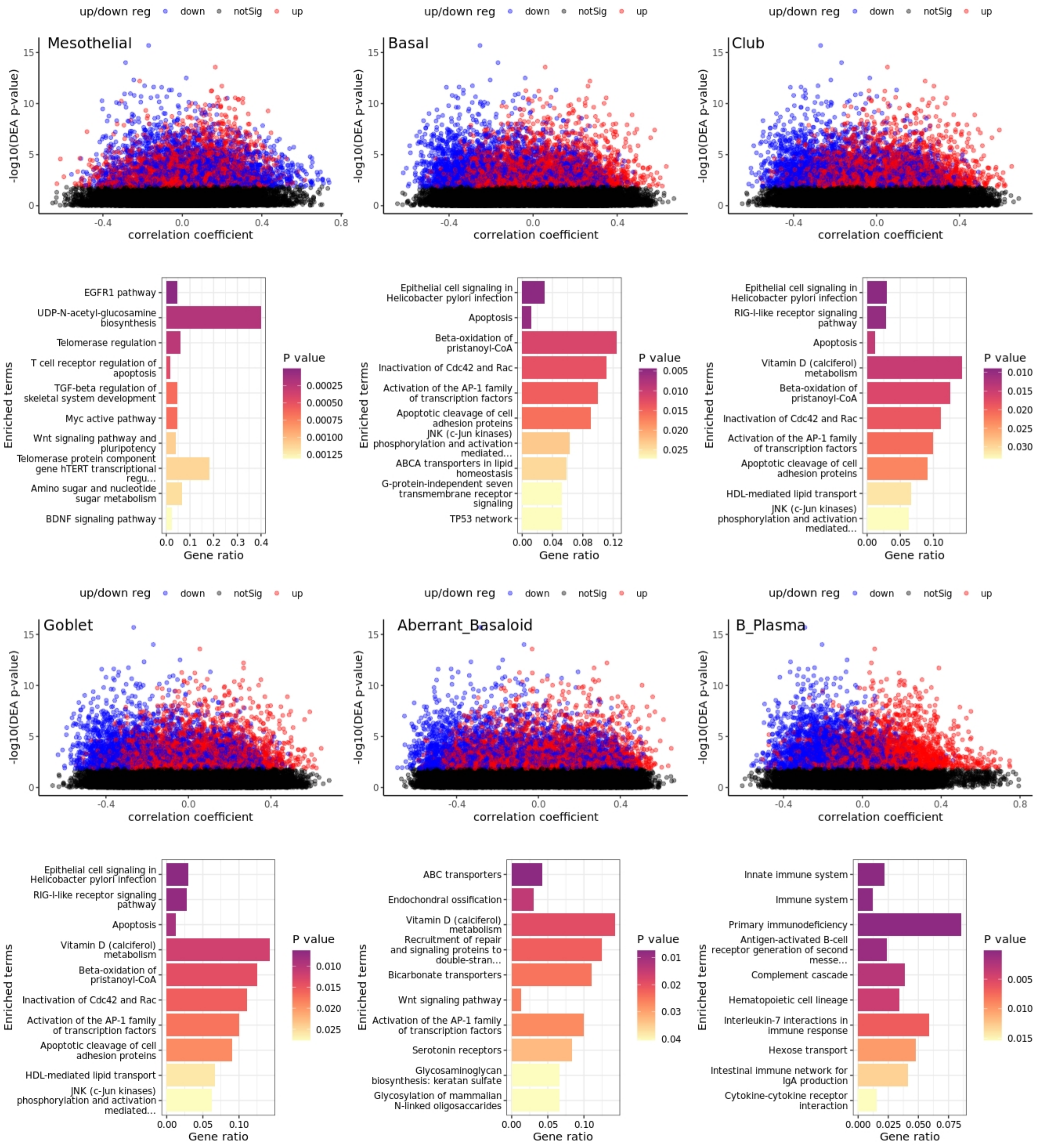


**Fig. S10 (cont'd).** Cell-type specific pathway analysis. $\log_{10}$p-value vs. correlation plots and pathways per cell type, after PC1-PC4 correction for GSE134692. Pathway result for each cell type is directly below its scatter plot.

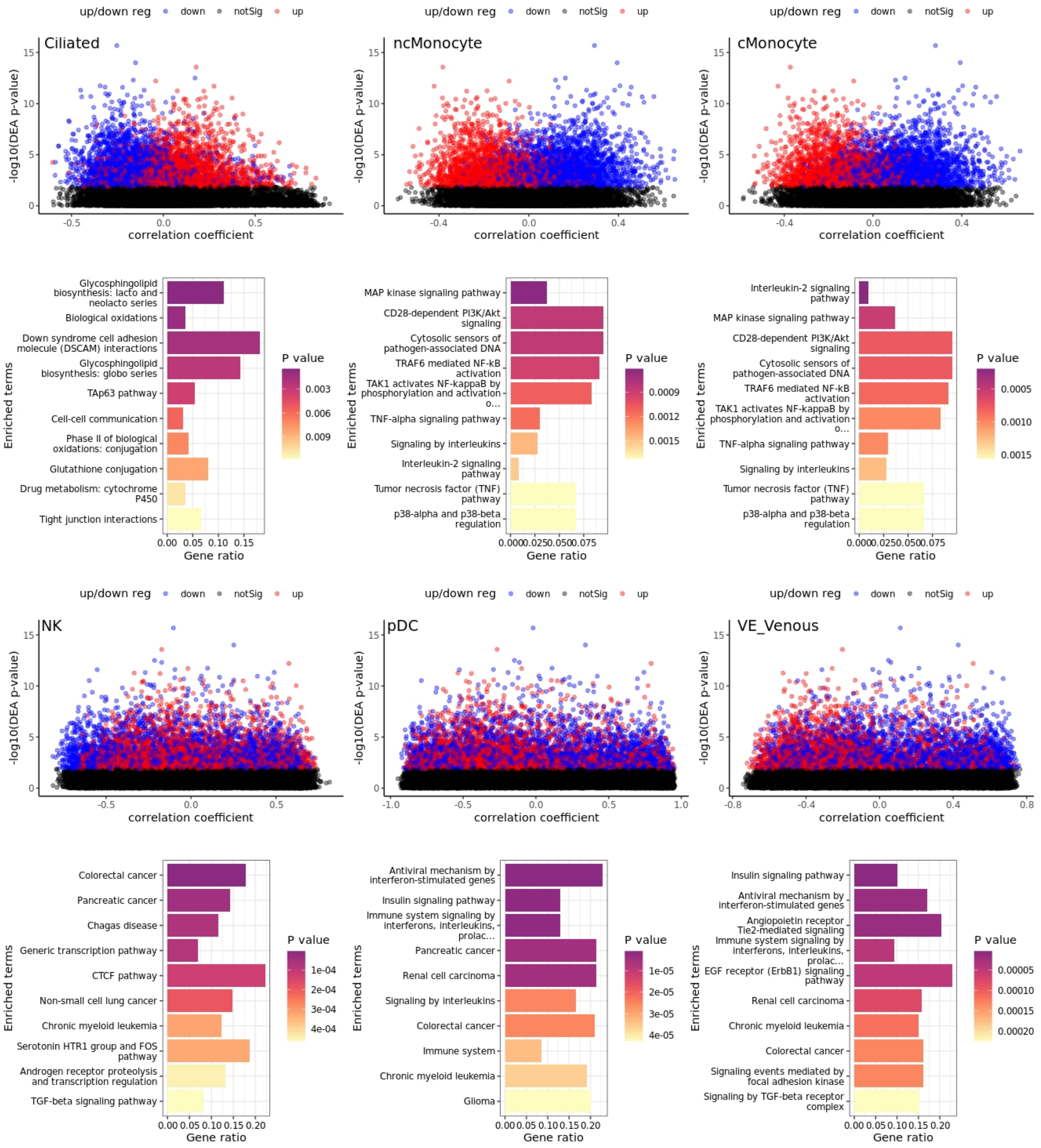


**Fig. S10 (cont'd).** Cell-type specific pathway analysis. $\log_{10}$p-value vs. correlation plots and pathways per cell type, after PC1-PC4 correction for GSE134692. Pathway result for each cell type is directly below its scatter plot.

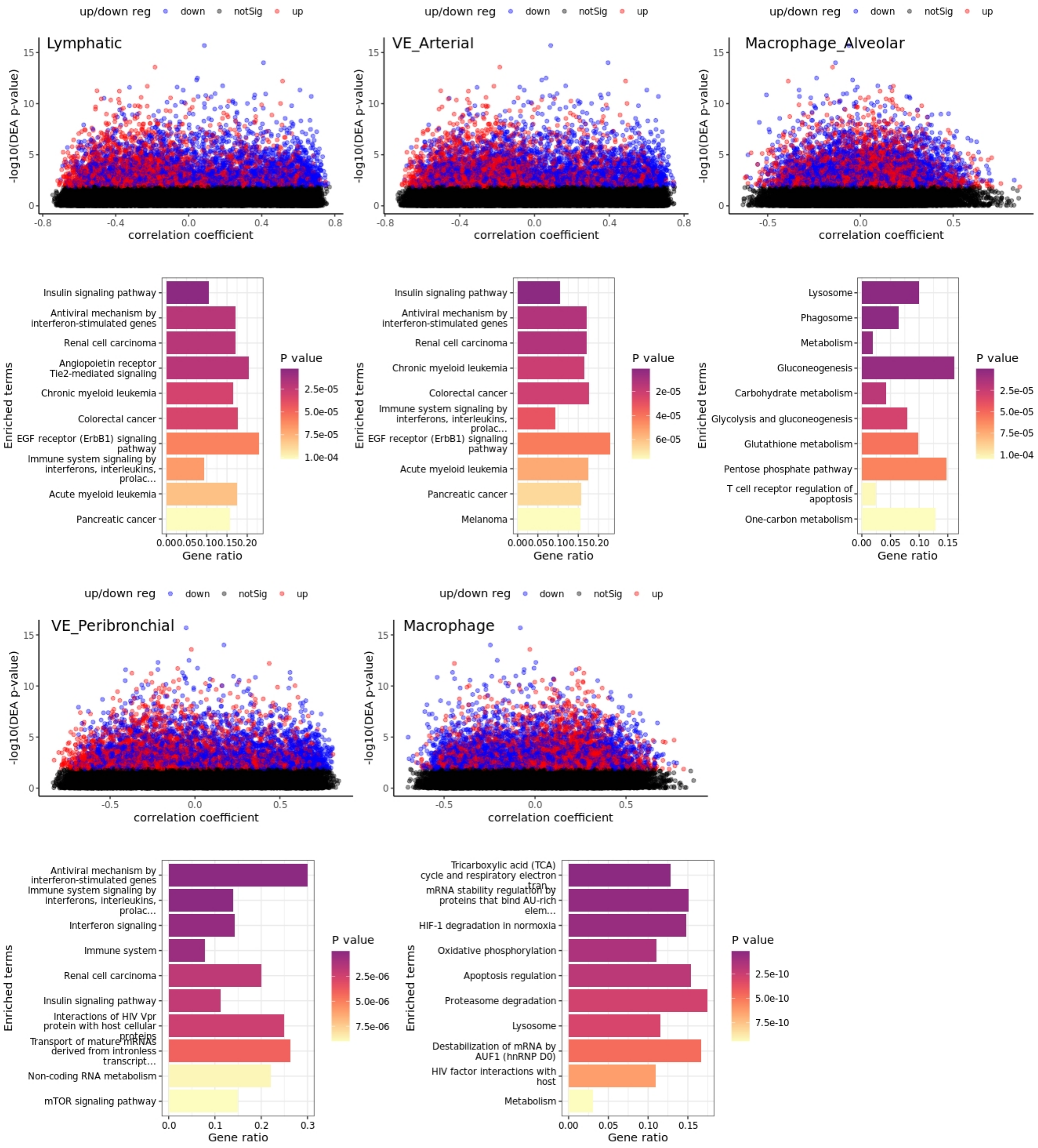


**Fig. S10 (cont'd).** Cell-type specific pathway analysis. $\log_{10}$p-value vs. correlation plots and pathways per cell type, after PC1-PC4 correction for GSE134692. Pathway result for each cell type is directly below its scatter plot.

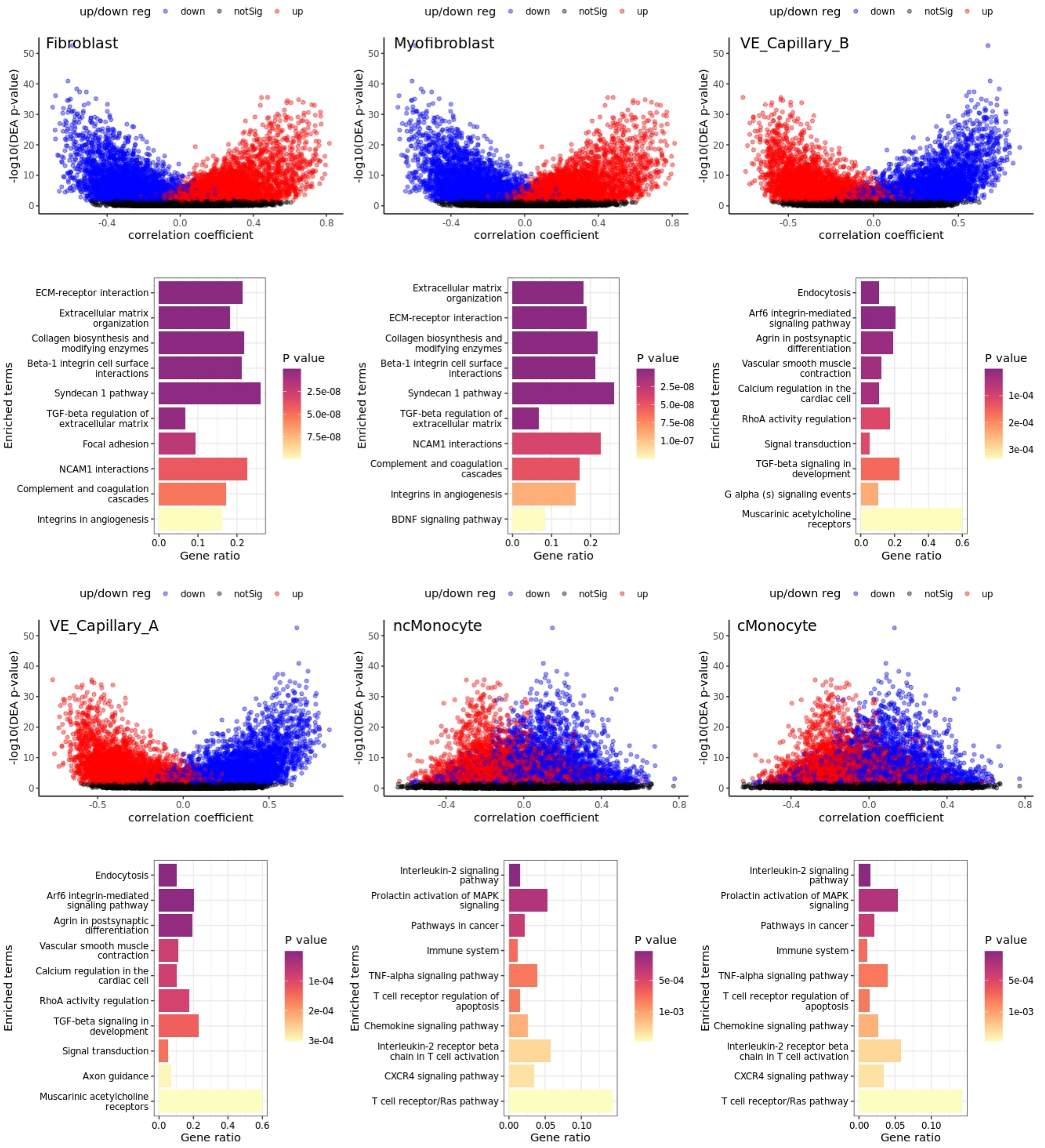


**Fig. S11.** Cell-type specific pathway analysis. $\log_{10}$p-value vs. correlation plots and pathways per cell type, after PC1 correction for GSE150910. Pathway result for each cell type is directly below its scatter plot.

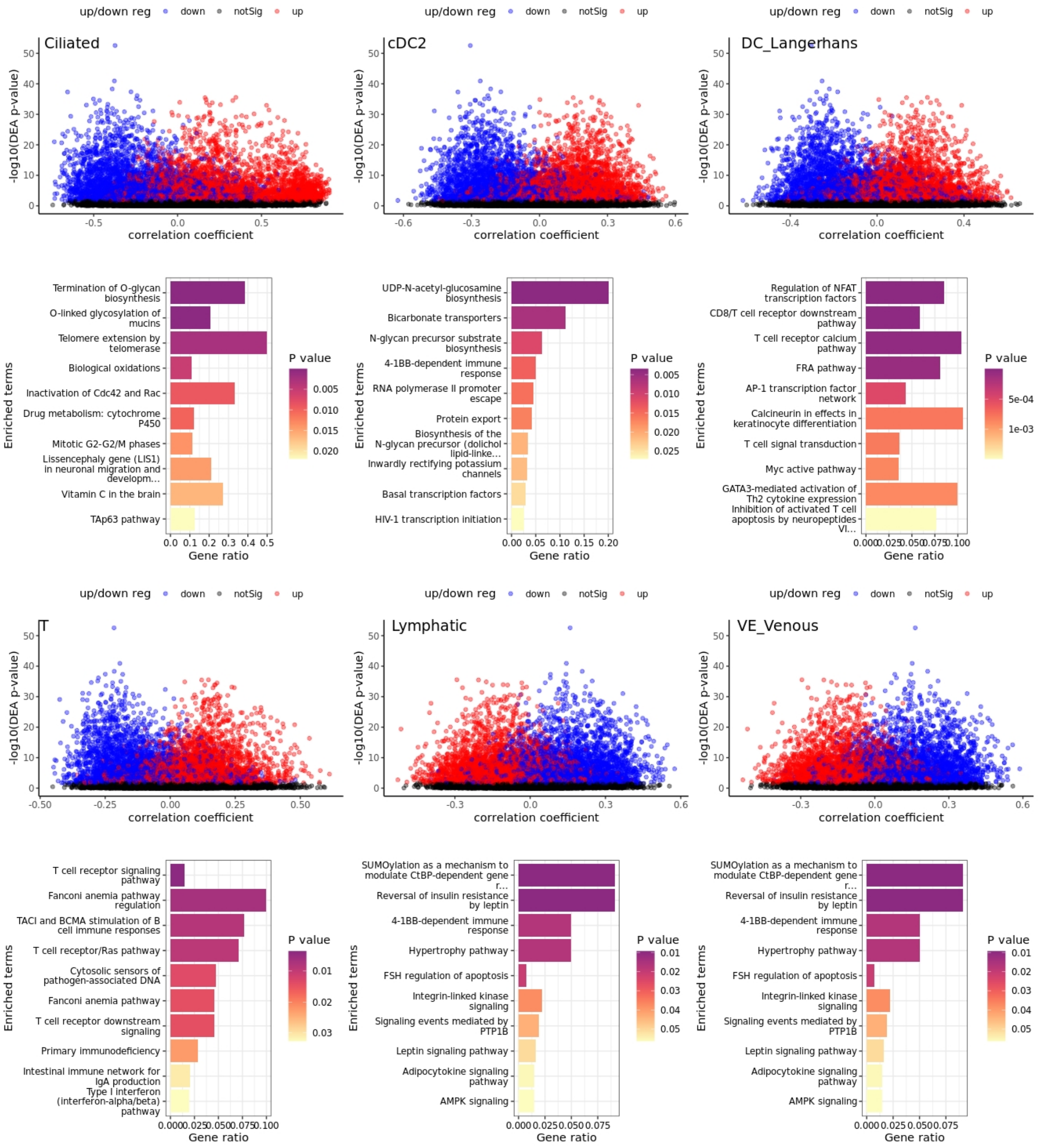


**Fig. S11 (cont'd).** Cell-type specific pathway analysis. $\log_{10}$p-value vs. correlation plots and pathways per cell type, after PC1 correction for GSE150910. Pathway result for each cell type is directly below its scatter plot.

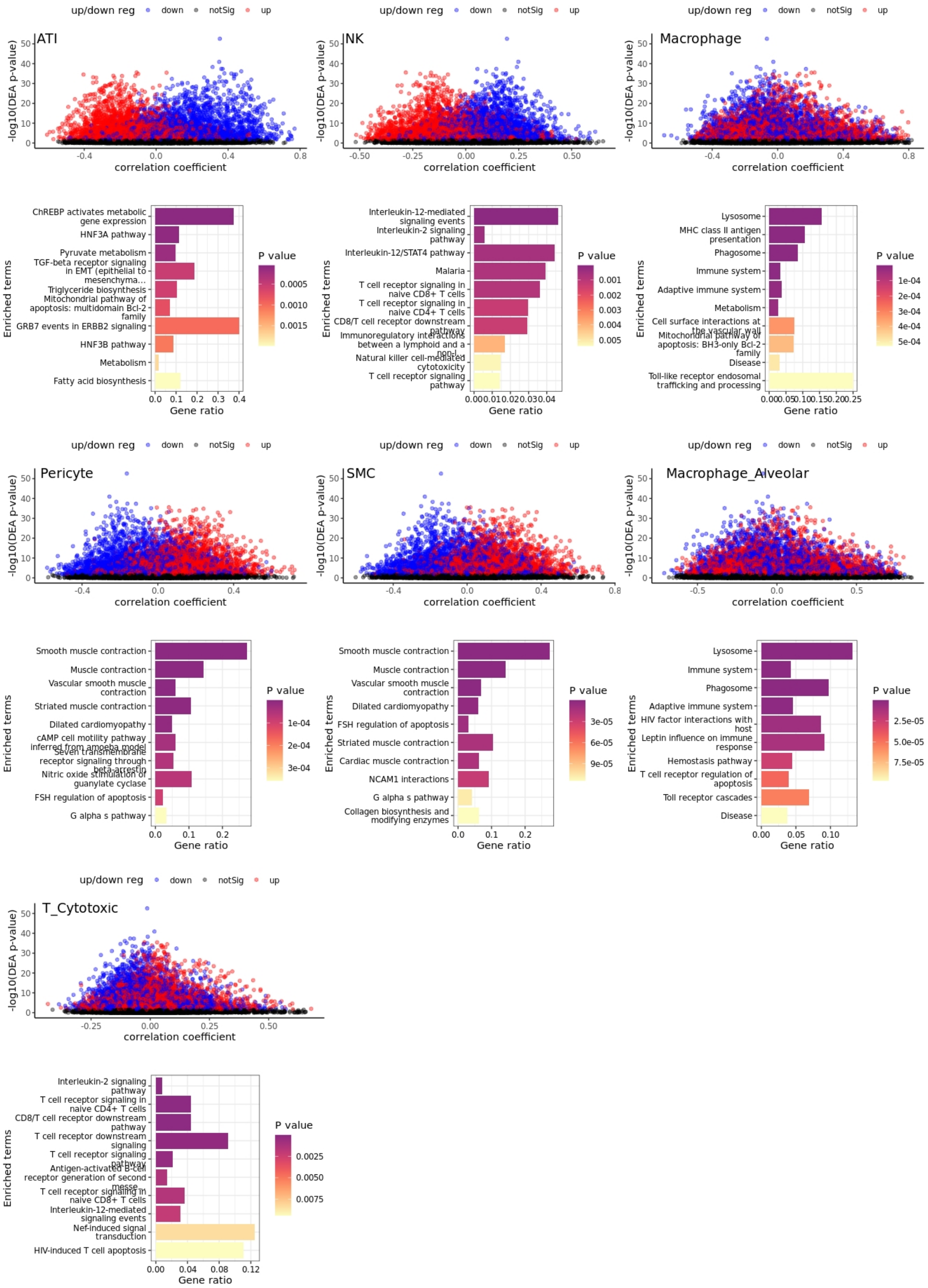


**Fig. S11 (cont'd).** Cell-type specific pathway analysis. $\log_{10}$p-value vs. correlation plots and pathways per cell type, after PC1 correction for GSE150910. Pathway result for each cell type is directly below its scatter plot.

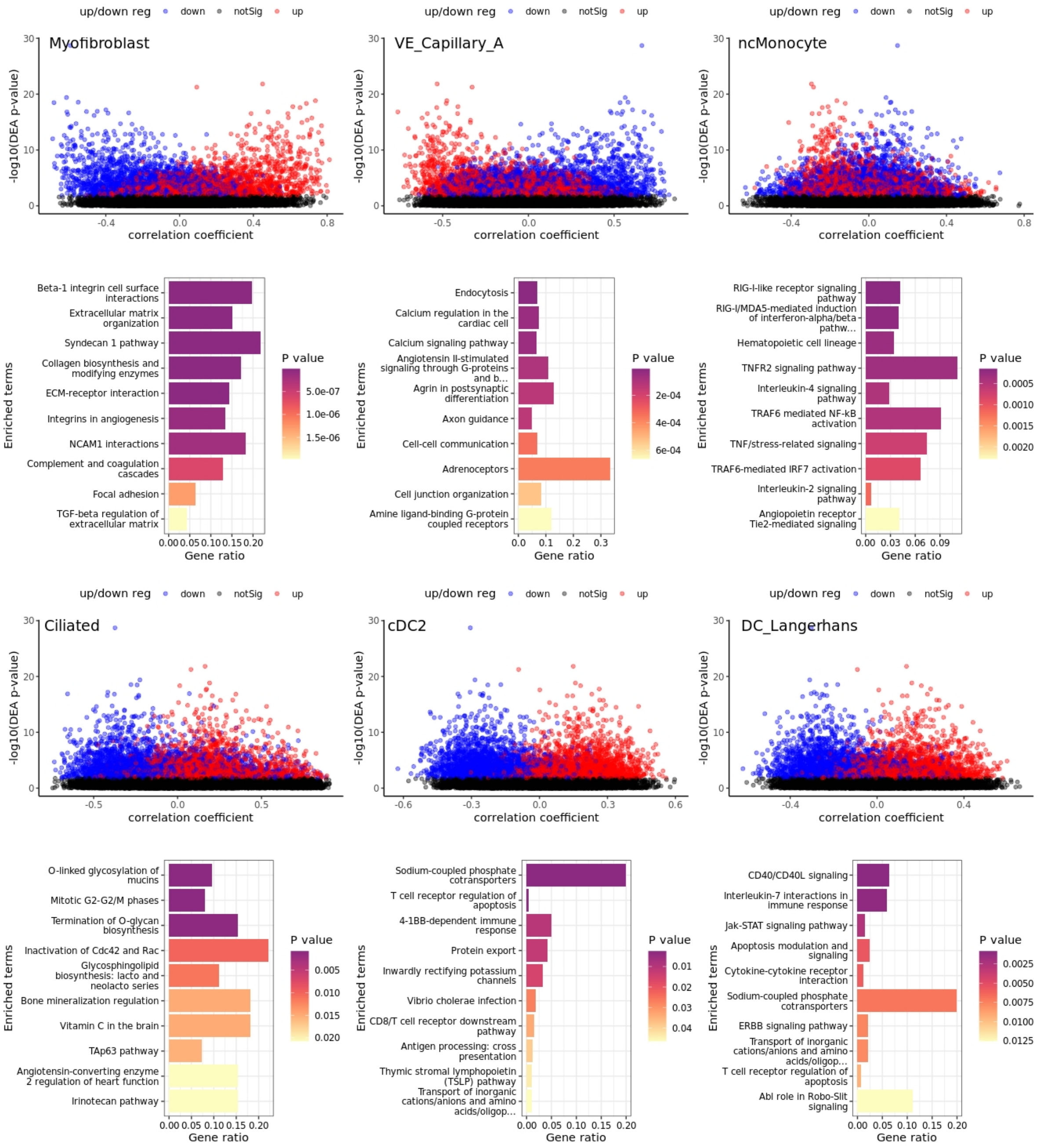


**Fig. S12.** Cell-type specific pathway analysis. $\log_{10}$p-value vs. correlation plots and pathways per cell type, after PC1-PC4 correction for GSE150910. Pathway result for each cell type is directly below its scatter plot.

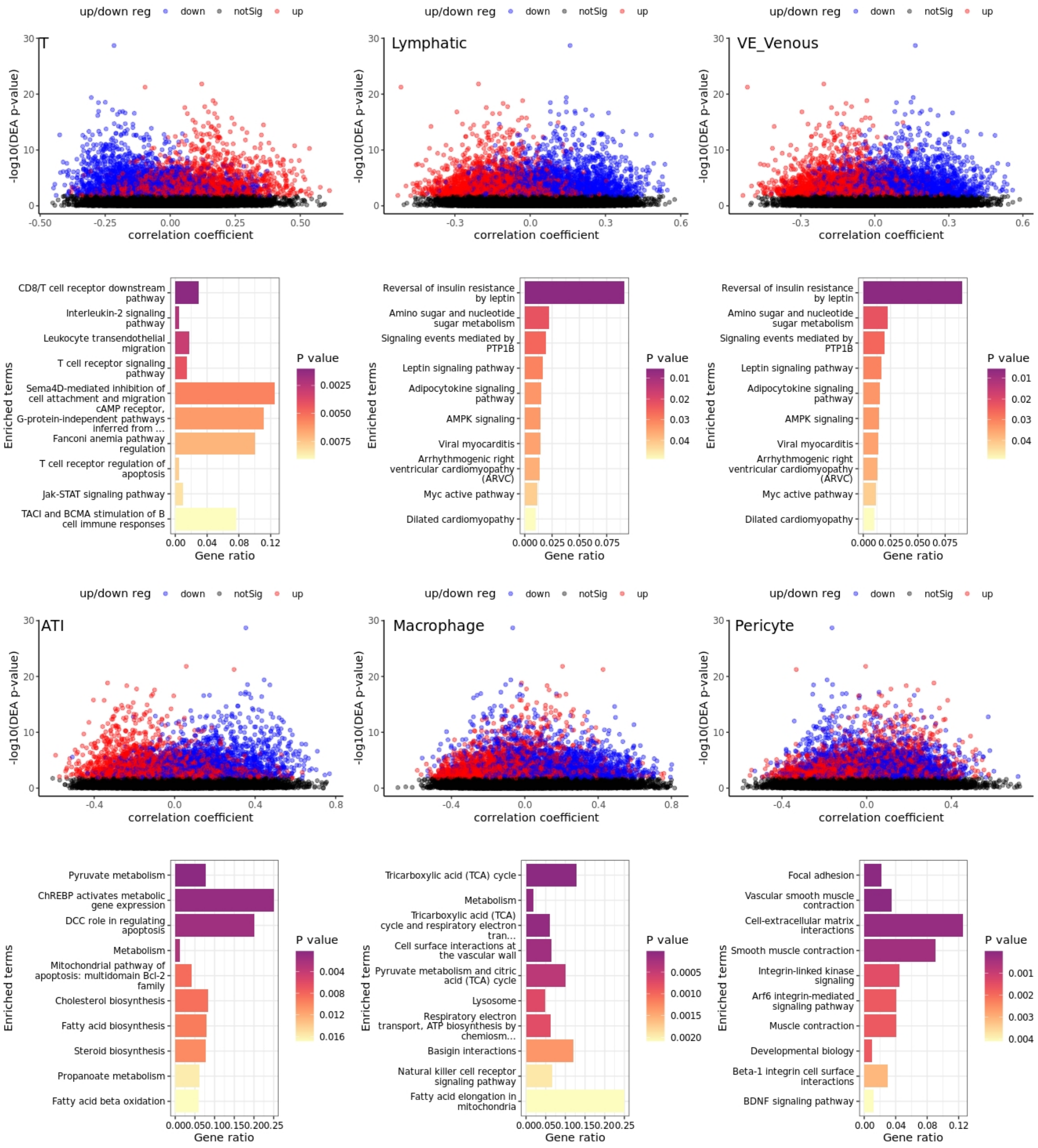


**Fig. S12 (cont'd).** Cell-type specific pathway analysis. $\log_{10}$p-value vs. correlation plots and pathways per cell type, after PC1-PC4 correction for GSE150910. Pathway result for each cell type is directly below its scatter plot.

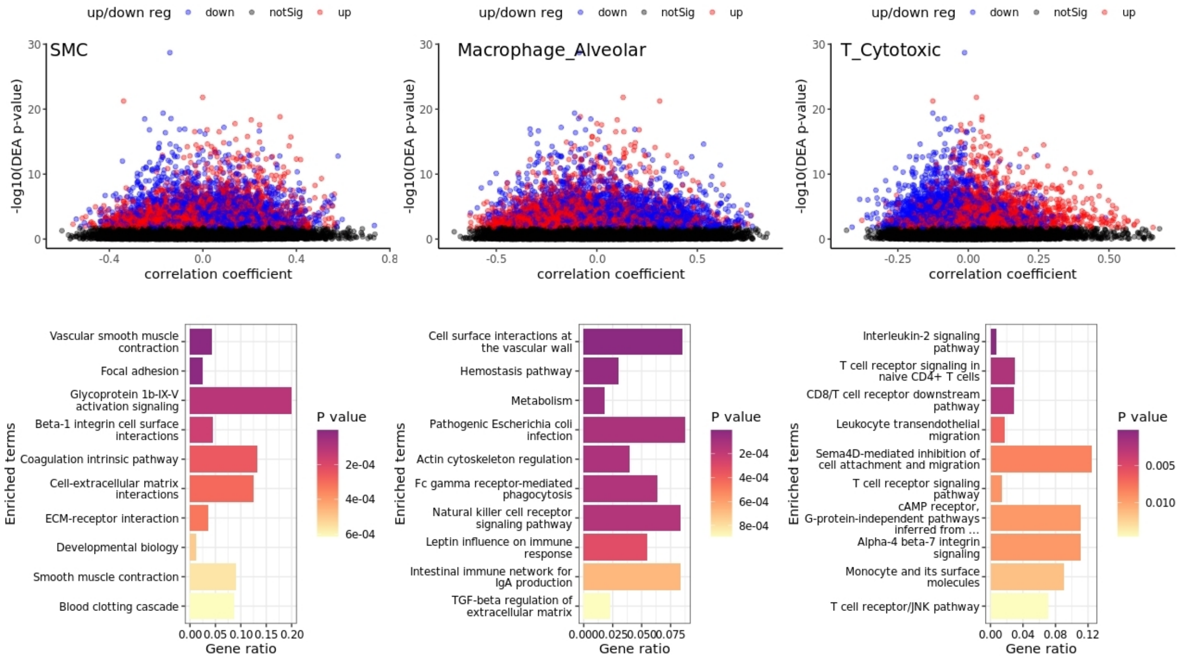


**Fig. S12 (cont'd).** Cell-type specific pathway analysis. $\log_{10}$p-value vs. correlation plots and pathways per cell type, after PC1-PC4 correction for GSE150910. Pathway result for each cell type is directly below its scatter plot.

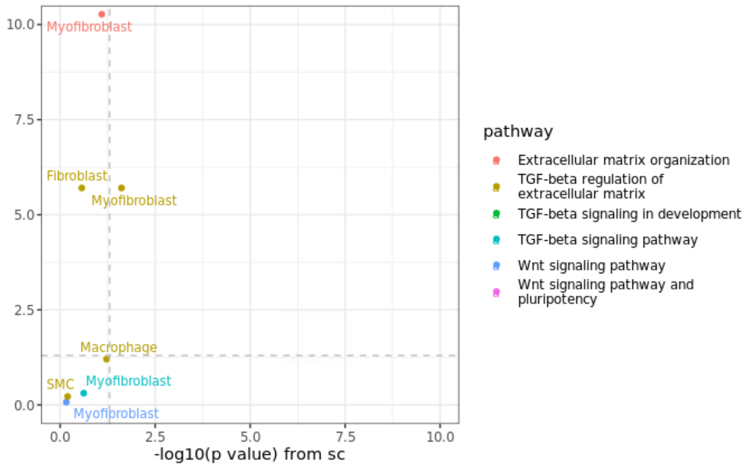


**Fig. S13.** Validation of cell type-specific pathways in bulk data GSE150910, after PC1-PC4 correction.
